# Control of Hox transcription factor concentration and cell-to-cell variability by an auto-regulatory switch

**DOI:** 10.1101/115220

**Authors:** Dimitrios K. Papadopoulos, Kassiani Skouloudaki, Ylva Engström, Lars Terenius, Rudolf Rigler, Christoph Zechner, Vladana Vukojević, Pavel Tomancak

## Abstract

The variability in transcription factor concentration among cells is an important developmental determinant, yet how variability is controlled remains poorly understood. Studies of variability have focused predominantly on monitoring mRNA production noise. Little information exists about transcription factor protein variability, since this requires the use of quantitative methods with single-molecule sensitivity. Using Fluorescence Correlation Spectroscopy (FCS), we characterized the concentration and variability of 14 endogenously tagged TFs in live *Drosophila* imaginal discs. For the Hox TF Antennapedia we investigated whether protein variability results from random stochastic events or is developmentally regulated. We found that Antennapedia transitioned from low concentration/high variability early, to high concentration/low variability later, in development. FCS and temporally resolved genetic studies uncovered that Antennapedia itself is necessary and sufficient to drive a developmental regulatory switch from auto-activation to auto-repression, thereby reducing variability. This switch is controlled by progressive changes in relative concentrations of preferentially activating and repressing Antennapedia isoforms, which bind chromatin with different affinities. Mathematical modelling demonstrated that the experimentally supported auto-regulatory circuit can explain the increase of Antennapedia concentration and suppression of variability over time.

## Introduction

In order to understand the mechanisms that control pattern formation and cell fate specification in developing organisms, the intranuclear concentration, DNA– binding kinetics and cell-to-cell variability of relevant transcription factors (TFs) need to be quantified. TF concentration variability at the tissue level is thought to arise from diverse processes, including mRNA transcription, translation and protein degradation. For example, transcription in a given tissue is noisy. Intrinsic noise is due to stochastic binding and interactions of proteins involved in transcriptional activation of a specific gene (Blake et al., 2003; Elowitz et al., 2002). Extrinsic noise arises from inter-cellular differences in abundance of the transcriptional and post-transcriptional machinery and affects the efficiency of transcriptional activation in general (Swain et al., 2002).

In undifferentiated tissue or cells, TF cell-to-cell variability can be the driving force for differentiation. For example, progressive establishment of a Nanog salt-and-pepper expression pattern leads to the formation of primitive endoderm in the mouse preimplantation embryo, whereas loss of the variability results in embryos lacking primitive endoderm entirely (Kang et al., 2013). In Drosophila, the Senseless (Sens) TF is required for the establishment of the proper number of sensory organ precursors in the ectodermal proneural cluster of cells and unequal Sens concentration among cells is required for their specification (Li et al., 2006). Moreover, variability in concentration (rather than its overall average concentration) of the Yan TF drives the transition of developing photoreceptor cells to a differentiated state during Drosophila eye development (Pelaez et al., 2015).

Conversely, in already differentiated tissue or cells, TF expression variability among cells may need to be counteracted to ensure homogeneity of gene expression patterns and robustness of commitment to a certain transcriptional regime. Examples are the Snail (Sna) TF, which is required for the invagination of the mesoderm during Drosophila gastrulation (Boettiger and Levine, 2013), or the Bicoid (Bcd) and Hunchback (Hb) TFs during early embryogenesis (Gregor et al., 2007a; Gregor et al., 2007b; Little et al., 2013). These studies have quantified the tolerable degrees of concentration variability that allow establishment of gene expression territories with remarkable precision in the developing embryo.

In addition, differential cell fates within the same developmental territory may be specified by TFs deploying different DNA-binding dynamics despite the existence of very similar concentrations (i.e. low variability). For example, studies on the Oct4 TF in early mouse embryos have shown that differential kinetic behavior of DNA binding, despite equal Oct4 concentration among blastomeres, ultimately dictates an early developmental bias towards lineage segregation (Kaur et al., 2013; Plachta et al., 2011).

So far, studies of gene expression variability have focused predominantly on monitoring the noise of mRNA production (Holloway et al., 2011; Holloway and Spirov, 2015; Little et al., 2013; Lucas et al., 2013; Pare et al., 2009). Little information exists about TF variability at the protein level within a tissue. Such studies require the use of quantitative methods with single-molecule sensitivity.

We have previously used fluorescence microscopy and Fluorescence Correlation Spectroscopy (FCS), to quantitatively characterize Hox TF interactions with nuclear DNA in living salivary gland cells (Papadopoulos et al., 2015; Vukojevic et al., 2010). FCS has also been instrumental for quantifying TF dynamics in living cells or tissue (Clark et al., 2016; Kaur et al., 2013; Lam et al., 2012; Mistri et al., 2015; Papadopoulos et al., 2015; Perez-Camps et al., 2016; Szaloki et al., 2015; Tiwari et al., 2013; Tsutsumi et al., 2016). However, in these studies, mobility has only been measured for overexpressed proteins. However, to understand TF behavior *in vivo,* proteins need to be quantified at endogenous levels (Lo et al., 2015).

In this study, we take advantage of the availability of fly toolkits, in which TFs have been endogenously tagged by different methodologies: fosmid (Baumgartner et al., 1996), BAC (deposition of lines of Rebecca Spokony and Kevin White to Flybase and the Bloomington Stock Center), FlyTrap (Buszczak et al., 2007; Kelso et al., 2004; Morin et al., 2001; Quinones-Coello et al., 2007) and MiMIC lines (Nagarkar-Jaiswal et al., 2015; Venken et al., 2011), to measure the intranuclear concentration of various TFs *in vivo* by FCS, and their cell-to-cell variability in fly imaginal discs. Imaginal discs are flat, single-layered epithelia comprised of small diploid cells and many TFs are expressed in defined regions within these tissues during development. Here, we found that Antp, a well-characterized homeotic selector gene, responsible for specification of thoracic identity in fly tissues, displays high cell-to-cell variability during early disc development and this variability was suppressed at later developmental stages. Through a combination of genetics, single-molecule measurements of TF dynamics by FCS and mathematical modeling, we uncovered a mechanism that controls Antp concentration and variability in developing discs.

## Results

### Characterization of average protein concentrations and cell-to-cell variability of *Drosophila* TFs

Average concentrations of TFs in neighboring nuclei of third instar imaginal discs were measured by FCS (Fig. 1A-J and Supplemental Fig. S1A-P). FCS is a non-invasive method with single molecule sensitivity, in which a confocal arrangement of optical elements is used to generate a small (sub-femtoliter) detection volume inside living cells, from which fluorescence is being detected (Fig. 1C,D; green ellipsoid). Fluorescent molecules diffuse through this observation volume, yielding fluorescence intensity fluctuations that are recorded over time by detectors with single-photon sensitivity (Fig. 1E). These fluctuations are subsequently subjected to temporal autocorrelation analysis, yielding temporal autocorrelation curves (henceforth referred to as FCS curves, Fig. 1F), which are then fitted with selected models to extract quantitative information about the dynamic processes underlying the generation of the recorded fluctuations. In the case of molecular movement of TFs (Supplement 1), information can be obtained regarding: a) the absolute TF concentrations (Fig. 1F), (b) TF dynamic properties, such as: diffusion times, differences in their interactions with chromatin and fractions of free-diffusing *versus* chromatin-bound TFs (Fig. 1G); and c) cell-to-cell TF concentration variability (Fig. 1H).

**Figure 1:**
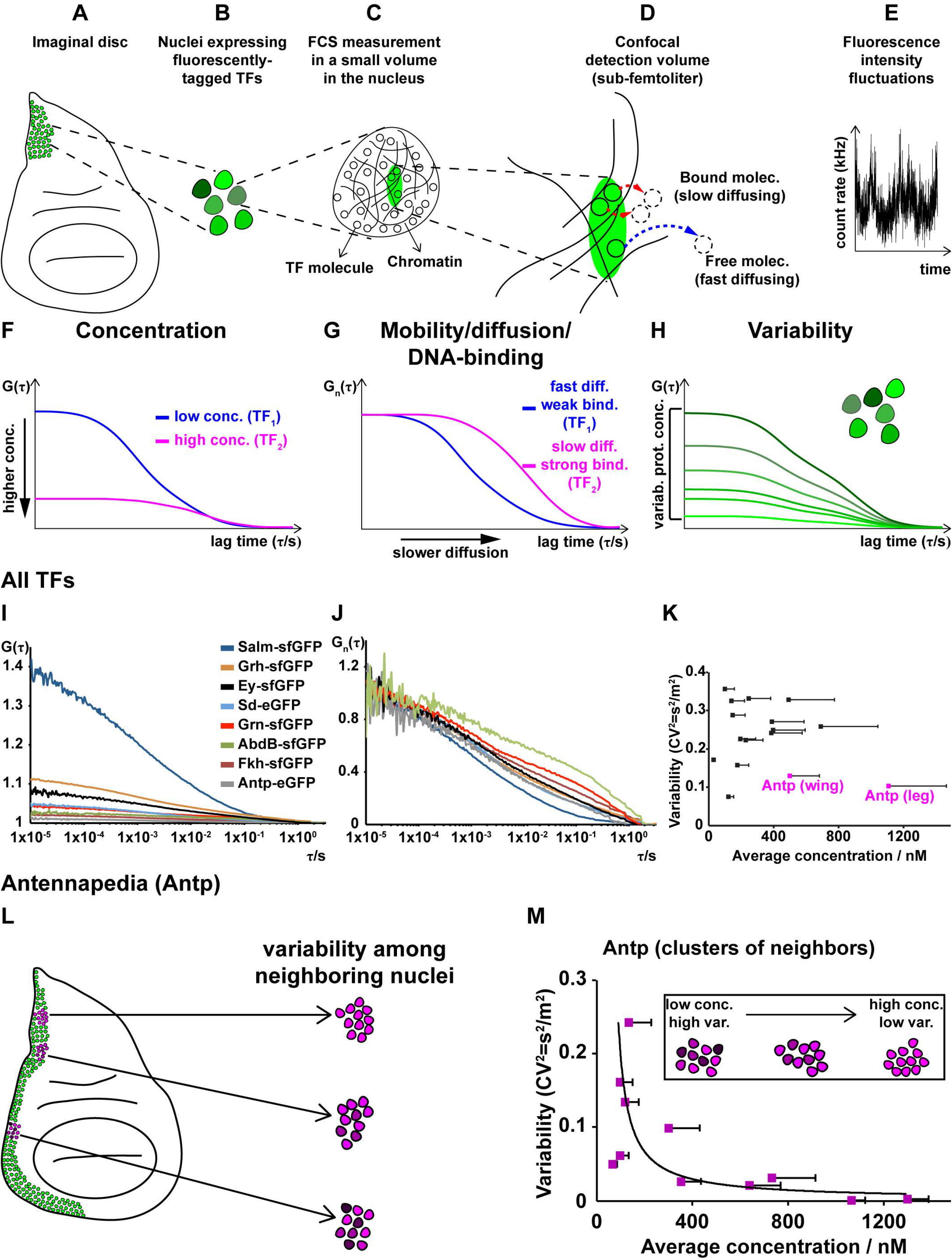
Concentration, DNA-binding dynamics and cell-to-cell protein concentration variability of 14 Drosophila TFs. (A-H) Workflow of the study of TFs by FCS (see Materials and Methods and Supplement 1). (I) Representative average FCS measurements of eight TFs. (J) FCS curves shown in (I), normalized to the same amplitude, *G_n_*(*τ*) = 1 at *τ* = 10 μ*s*. (K) Variability of the 14 TFs as a function of concentration. (L) Variability in concentration of endogenous Antp in the wing disc. (M) Variability of Antp concentration in clusters of neighboring cell nuclei as a function of concentrations. Error bars in (K) and (N) represent 1 standard deviation.

We selected 14 TF based on the availability of homozygous, endogenously tagged transgenes and generation of robust fluorescence in distinct patterns in various imaginal discs. For the 14 TFs, we measured average concentrations ranging about two orders of magnitude among different TFs, from ~30 *nM* to ~1.1 *μM*(~400 to 15500 molecules per nucleus, respectively) (Fig. 1I, Supplemental Fig. S1A-Q and Supplement 2). We also obtained various diffusion times and fractions of slow and fast diffusing TF molecules (Fig. 1J), indicating differential mobility and degree of DNA-binding among different TFs (Vukojevic et al., 2010). Comparison of the y-axis amplitudes at the zero lag time of the FCS curves, which are inversely proportional to the concentration of fluorescent molecules (Fig. 1F), gives information about concentration variability (heterogeneity) among different cell nuclei, i.e. reflects heterogeneity of protein concentration at the tissue level (Fig. 1H). For all 14 TFs studied, the variability, expressed as the variance over the mean squared, 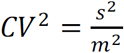 was determined to be in the range 7 – 37% (Fig. 1K and Supplemental Fig. S1Q).

In biological systems, the Fano factor, expressed as the variance over the mean (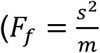, in concentration units), is a commonly used index to quantify variability. It has been proposed that Fano factor values that increase with average concentrations indicate that the underlying transcriptional processes cannot be sufficiently explained by a simple one-step promoter configuration with purely intrinsic Poissonian noise and that extrinsic noise is likely to contribute significantly to the overall variability (Newman et al., 2006; Schwanhausser et al., 2011; Taniguchi et al., 2010). For all TFs measured, Fano factor values from 0 to 20 were obtained (Supplemental Fig. S1R), in line with Fano factor values of other TFs determined previously to lie between 0 and 30 (Sanchez et al., 2011). Moreover, the majority of TFs examined show Fano factor values, F_f_ > 1, suggesting that transcriptional bursting is likely to be a significant source of the observed cell-to-cell variability. For Eyeless, the Fano factor value F_f_ ≈ 1 suggests that the transcription of this gene is likely to be a continuous process that occurs with a constant probability and according to the statistics of a Poisson process. In contrast, Salm, which was the only TF characterized by F_f_ < 1, exhibits a pattern of occurrence that is more regular than the randomness associated with a Poissonian process. We used this dataset as a starting point for studying the control of variability during imaginal disc development.

The average concentration and variability of the investigated TFs showed no obvious interdependence (Fig. 1K), suggesting that if variability is controlled, there is not one control mechanism that is common to all investigated TFs. Among the studied TFs, the Hox protein Antennapedia (Antp), showed low variability (*CV*^2^ < 0.2) in high average concentrations, in particular in the leg disc (Fig. 1K). Since low variability at the tissue level is likely to be achieved through regulatory mechanisms, we investigated Antp variability further by FCS. Because FCS performs best at low to moderate expression levels (Supplement 1), we performed this analysis in the wing disc where expression levels are lower than in the leg disc (Fig. 1K,L). We first established that the observed fluorescence intensity fluctuations were caused by diffusion of TF molecules through the detection confocal volume (Supplemental Fig. S2 and Supplement 1). FCS showed that different clusters of neighboring cells along the Antp expression domain in the wing disc display different average expression levels (Fig. 1L). Moreover, FCS showed that Antp cell-to-cell variability decreased with increasing Antp concentration (Fig. 1M) whereas the Fano factor increased (Supplemental Fig. S1R). Such behavior is indicative of complex transcriptional regulatory processes (Franz et al., 2011; Smolander et al., 2011) that we further investigated using the powerful *Drosophila* genetic toolkit.

### Control of Antp concentration by transcriptional auto-regulation

One mechanism by which genes control their expression level variability is auto-regulation (Becskei and Serrano, 2000; Dublanche et al., 2006; Gronlund et al., 2013; Nevozhay et al., 2009; Shimoga et al., 2013; Thattai and van Oudenaarden, 2001). To test whether Antp can regulate its own protein levels, we monitored the concentration of endogenous Antp protein upon overexpression of *Antp* from a transgene. To distinguish between overexpressed and endogenous protein, we used synthetic Antp (SynthAntp) transgenes fused to eGFP (SynthAntp-eGFP). These transgenes encode the Antp homeodomain, the conserved YPWM motif and the C terminus (but lack the long and non-conserved N terminus of the protein, against which widely used Antp antibodies have been raised) and they harbor Antp-specific homeotic function (Papadopoulos et al., 2011). Clonal overexpression of *SynthAntp-eGFP* in the wing disc notum (Fig. 2A-B’ and controls in Supplemental Fig. S3D,D’) repressed the endogenous Antp protein, indicating that Antp is indeed able to regulate its own protein levels.

**Figure 2:**
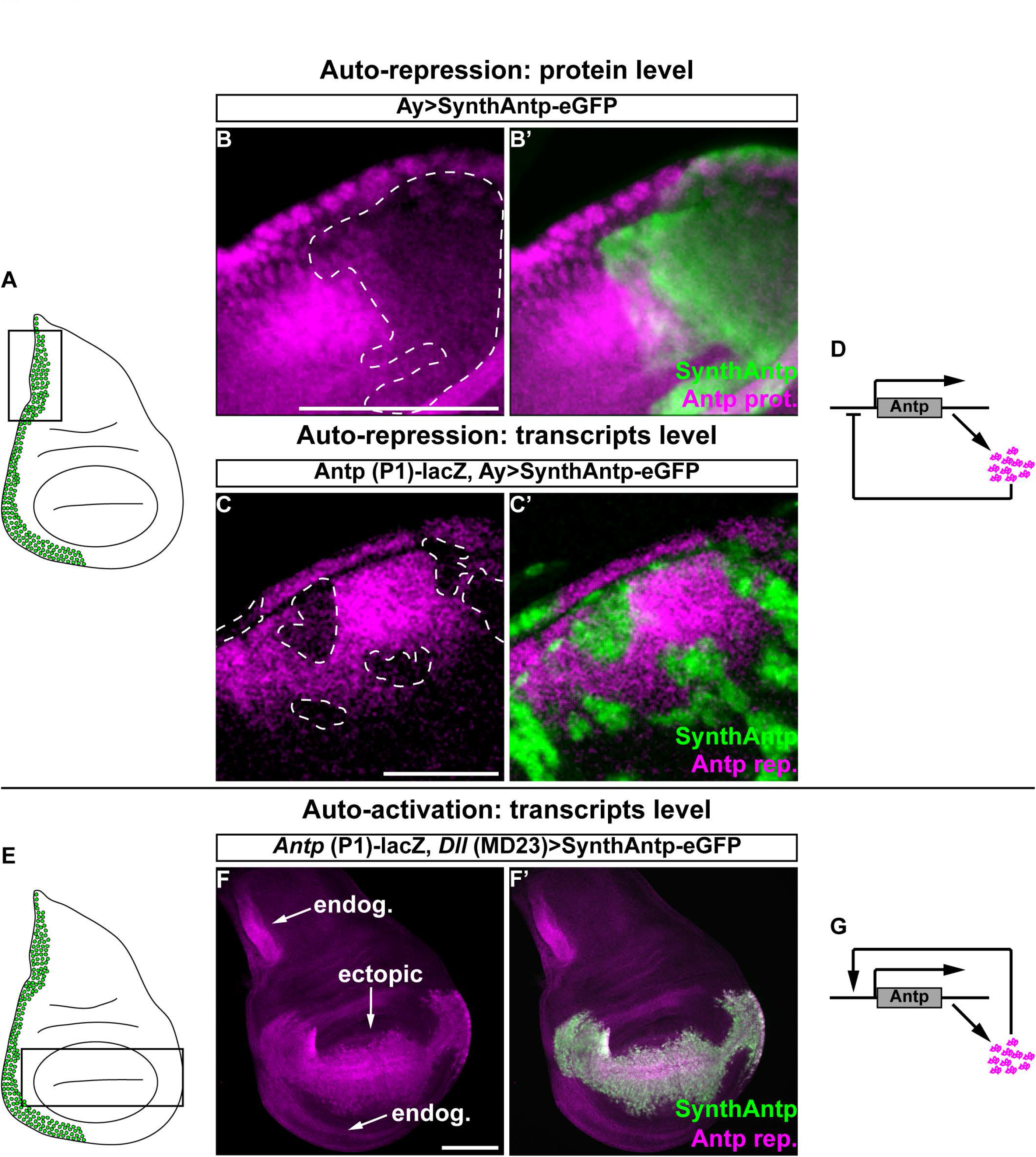
Antp activates and represses its own transcription. (A) Schematic of the wing disc used for analysis. (B-B’) Clonal overexpression of a *SynthAntp-eGFP* construct. Dashed line in (B) shows a clone in the Antp expression domain. (C-C’) Transcriptional auto-repression of Antp using the *Antp* P1-*lacZ.* (D) Schematic of Antp transcriptional auto-repression. (E) Schematic of the wing disc used for analysis. (FF’) Ectopic induction of *Antp* P1-*lacZ* by expression of *SynthAntp-eGFP* using *Dll-*Gal4 (MD23). (G) Schematic of Antp auto-activation. Scale bars denote 100 *μm*.

Since Antp is a TF, we next asked whether the auto-repression indeed occurs at the transcriptional level. The *Antp* locus is subject to complex transcriptional regulation, involving a distal and a proximal promoter (P1 and P2 promoters, respectively), spanning more than 100 kb of regulatory sequences. We established that the P1 promoter (rather than the P2 promoter) is predominantly required to drive expression of Antp in the wing disc notum (Supplemental Fig. S3A-C’), in line with previous observations ((Engstrom et al., 1992; Jorgensen and Garber, 1987; Zink et al., 1991) and Materials and Methods). Moreover, mitotic recombination experiments in regions of the wing disc unique to P2 transcription have shown no function of the P2 promoter transcripts in wing disc development (Abbott and Kaufman, 1986). Thus, the P1 Antp reporter serves as a suitable reporter of the *Antp* locus transcriptional activity in this context.

Clonal overexpression of SynthAntp-eGFP in the wing disc repressed the *Antp* P1 transcriptional reporter (Fig. 2C,C’ and controls in Supplemental Fig. S3E,E’). To rule out putative dominant negative activity of the small SynthAntp-eGFP peptide, we also performed these experiments with the full-length Antp protein (Supplemental Fig. S3F,F’). We conclude that the Antp protein is able to repress its own transcription from the P1 promoter, suggesting a possible mechanism of suppressing cell-to-cell variability of *Antp* expression levels (Fig. 2D).

In the course of these experiments, we noticed that ectopic overexpression of *SynthAntp-eGFP* or the full-length Antp protein from the *Distal-less* (*Dll*) (MD23) enhancer resulted in activation of the *Antp* P1 reporter in distal compartments of the wing disc, such as the wing pouch, where Antp is normally not detected (Fig. 2E-F’ and controls in Supplemental Fig. S3G-H’). This suggests that next to its auto-repressing function, Antp is also capable of activating its own transcription (Fig. 2G).

To exclude that the auto-activation and repression of Antp are artifacts of overexpression, we used FCS to measure the concentration of Antp triggered by different Gal4-drivers (Supplemental Fig. S4A-E). We observed indistinguishable DNA-binding behavior by FCS, not only across the whole concentration range examined (Supplemental Fig. S4F), but also between endogenous and overexpressed Antp (Supplemental Fig. S5A,B). Importantly, the auto-activating and auto-repressing capacity of Antp was preserved even with the weak Gal4-driver *69B* (Supplemental Fig. S4K,L) that triggered concentrations of Antp lower than its normal concentration in the leg disc (473 *nM* versus 1110 *nM*), indicating that auto-activation and auto-repression of Antp take place at endogenous protein concentrations.

We conclude that Antp is able to repress and activate its own transcription (Fig. 2D,G) and hypothesize that this auto-regulatory circuit sets the correct concentration of Antp protein in imaginal discs.

### A temporal switch controls the transition of *Antp* from a state of auto-activation to a state of auto-repression

To further investigate the mechanism by which the Antp auto-regulatory circuit sets the precise Antp expression levels, we next asked whether the seemingly opposing auto-regulatory activities of Antp are separated in time during development. To that end, we induced gain-of-function clones of full-length untagged *Antp* either at 26 h (first larval instar – henceforth referred to as “early” stage) or at 60 h (late second larval instar – henceforth referred to as “late” stage) of development and analyzed the clones in late third instar wing imaginal discs (Fig. 3). As a pre-requisite for this analysis, we established that the Antp-eGFP homozygous viable MiMIC allele recapitulates the endogenous Antp pattern in the embryo and all thoracic imaginal discs and therefore can be used to monitor endogenous Antp protein (Supplemental Fig. S6). Clonal induction of full-length untagged *Antp* in early development triggered strong auto-activation of Antp-eGFP (Fig. 3A,B,B’ and controls in Supplemental Fig. S7A-C’). As before, we confirmed that early auto-activation of Antp is transcriptional and similar for both full-length and SynthAntp proteins (Supplemental Fig. S7D-E’ and controls in F-G’). Early auto-activation was further supported by a loss-of-function experiment, where *RNAi*-mediated early knockdown of Antp resulted in downregulation of the Antp reporter (Fig. 3C,C’ and controls in Supplemental Fig. S7H,H’). The loss and gain-of-function analysis together suggest that during early disc development Antp is required for sustaining its own expression.

**Figure 3:**
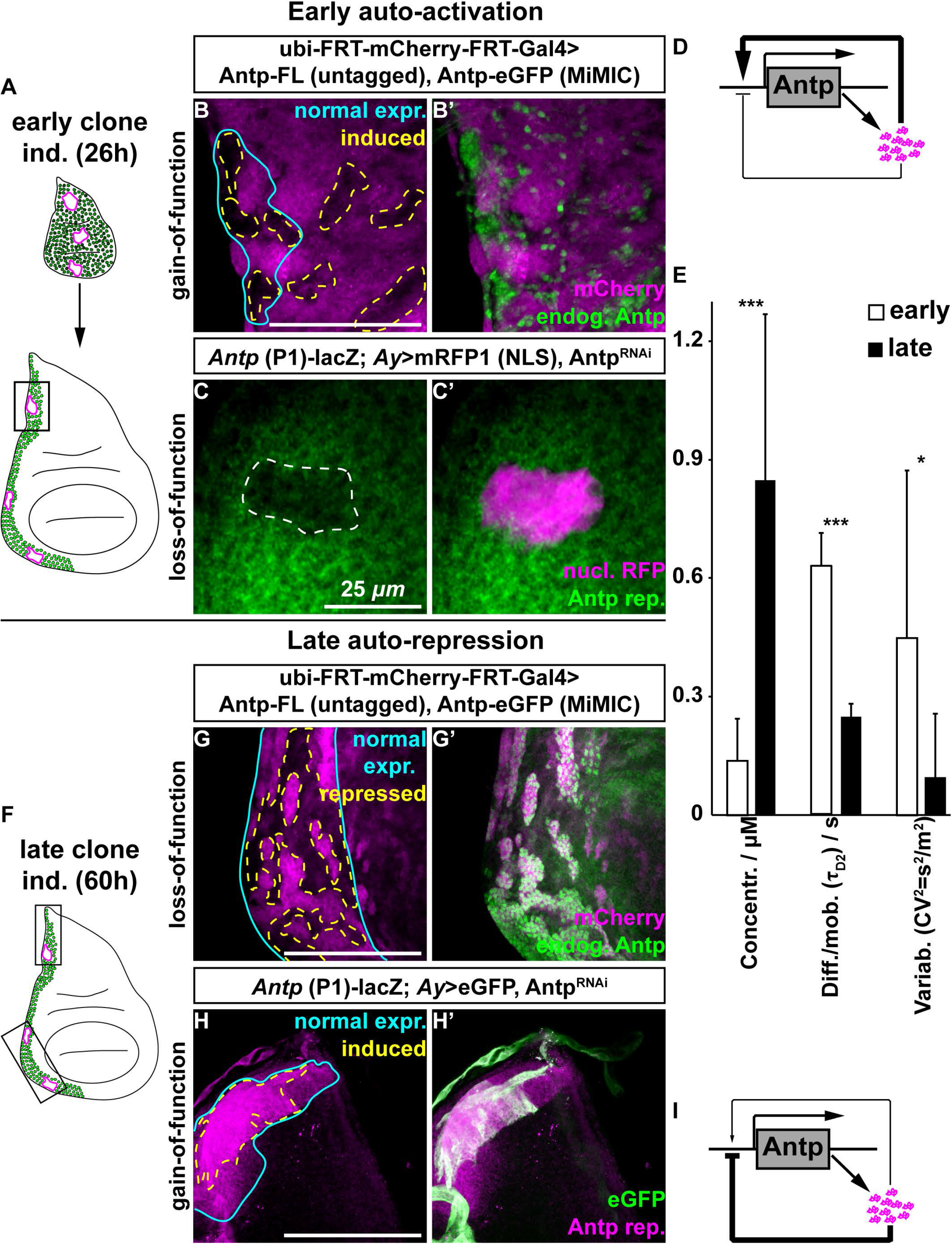
Antp switches from transcriptional auto-activation to auto-repression. (A and F) Schematic representations of the experimental setup (see Materials and Methods). Black rectangles enclose the regions of clonal analysis used. (B-B’) Endogenous Antp in early-induced clones, marked by the absence of mCherry (dashed lines in (B)). (C-C’) *Antp* P1 transcription in early *Antp RNAi* knockdown clones (dashed line in (C), marked by nuclear mRFP1. (D) Updated Antp autoactivation model. (E) Concentration, DNA–binding and variability studied by FCS at second instar leg and wing discs (FCS analysis in Supplemental Fig. S8). (G-G’) Endogenous Antp in late-induced clones, marked by the absence of mCherry (dashed lines in (G)). (H-H’) *Antp* P1 transcription in early *Antp RNAi* knockdown clones (dashed line in (H)), marked by nuclear mRFP1. (I) Updated Antp auto-repression model. Cyan lines in (B, G, H) outlines the region of highest endogenous Antp expression.

In contrast, clonal induction during the late second instar stage (Fig. 3F) repressed Antp-eGFP (Fig. 3G,G’) and, reciprocally, the clonal knockdown by *RNAi* triggered auto-activation of Antp transcription (Fig. 3H,H’). Hence, in contrast to early development, Antp represses its own expression in third instar discs.

While the gain-of-function experiments show that Antp is sufficient to execute auto-regulation, loss-of-function analysis indicates that it is also necessary for both repression and activation at the transcriptional level.

Together, these results revealed the existence of a switch in Antp auto-regulatory capacity on its own transcription during development. Starting from a preferentially auto-activating state early in development (Fig. 3D), Antp changes to an auto-inhibitory mode at later developmental stages (Fig. 3I).

### During development Antp switches from a low-concentration/high-variability to a high-concentration/low-variability state

If the *Antp* auto-repressive state limits the variability of Antp protein concentration among neighboring cells late in development, we expected that the variability would be higher during earlier stages, when auto-repression does not operate. We, therefore, used FCS to characterize the endogenous expression levels and cell-to-cell variability of Antp concentration in nuclei of second instar wing and leg discs. We observed significantly lower average concentrations of Antp protein in second versus third instar wing and leg discs and the inverse was true for concentration variability (Fig. 3E and Supplemental Fig. S8A,A’,C), indicating that the developmental increase in concentration is accompanied by suppression of concentration variability. In addition, FCS revealed a notable change in Antp characteristic decay times (signifying molecular diffusion, limited by chromatin-binding) at early versus late stages (Supplemental Fig. S8B). This behavior indicates that endogenous Antp is initially moving fast in the nucleus, as it undergoes considerably fewer interactions with chromatin compared to later stages where its interactions with chromatin are more frequent and longer lasting.

Taken together, our FCS measurements show that *Antp* is expressed at relatively low and highly variable levels in early developing discs, when genetic evidence indicates auto-activation capacity on its own transcription. Later in development, when Antp has reached a state of higher average concentrations, auto-repression kicks in, resulting in considerably lower variability among neighboring cells.

### Dynamic control of Antp auto-regulation by different Antp isoforms

The changing binding behavior of Antp on chromatin from second to third instar discs and the developmental transition from an auto-activating to an auto-repressing state suggested a causal relationship between the two phenomena. We, therefore, sought to identify molecular mechanisms that could link the observed changes in Antp chromatin-binding to Antp auto-activation and repression. It is well established that the Antp mRNA contains an alternative splice site in exon 7 immediately upstream of the homeobox-containing exon 8, and generates Antp isoforms differing in as little as 4 amino acids in the linker between the YPWM motif (a cofactor-interacting motif) and the homeodomain (Fig. 4A) (Stroeher et al., 1988). Our previous observation that long linker isoforms favor transcriptional activation of Antp target genes, whereas short linker isoforms favor repression of Antp targets (Papadopoulos et al., 2011), prompted us to examine whether the linker length is also responsible for differences in auto-regulation.

**Figure 4:**
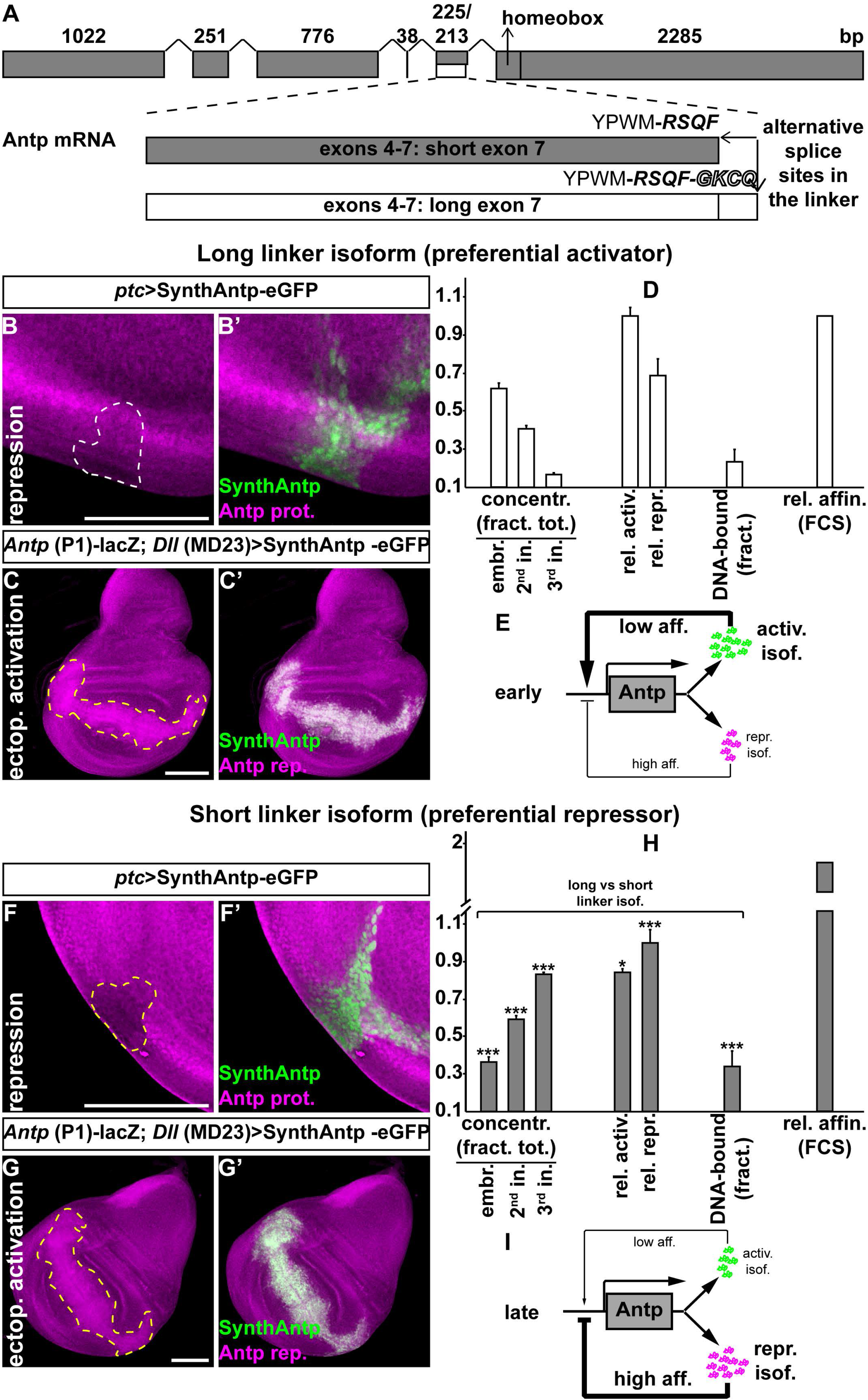
Antp auto-activation and auto-repression relies on Antp isoforms with different binding affinities to chromatin. (A) Schematic of the Antp mRNA, generated from the P1 promoter. Exons are represented by grey boxes. (B-B’) *Antp* auto-repression at the proximal wing disc (white dashed line). (C-C’) *Antp* autoactivation at the proximal wing disc (yellow dashed line). (D) Abundance of long linker isoform (Materials and Methods); auto-activation and auto-repression efficiencies (Materials and Methods); DNA-bound fractions, measured by FCS (<u>Supplemental Figure S10</u>); and relative affinity of binding to chromatin, measured by FCS (<u>Supplemental Figure S10</u>) for comparison with (H). (E) Updated model of *Antp* auto-regulation. (F-G’) Similar to (B-C’) for the short linker isoform. (H) Similar to (D) for comparison. (I) Updated qualitative model representation of Antp repression as in (E).

Ectopic expression of SynthAntp-eGFP peptides featuring a long linker displayed significantly weaker repression capacity on endogenous Antp, as compared to their short linker counterparts (Fig. 4B,B’,F,F’ and quantified in D,H, see also Materials and Methods). We confirmed that, also in this case, the repression was at the transcriptional level (Supplemental Fig. S9I-J’). Inversely, long linker Antp isoforms exhibited stronger activation of Antp reporter, as compared to short linker isoforms (Fig. 4C,C’,G,G’ and quantified in D,H; see also Materials and Methods). We, additionally, validated that short linker isoforms encoded by full-length or SynthAntp cDNAs behaved as weaker auto-activating and stronger auto-repressing Antp species in all our previous experiments using the endogenous Antp protein and the P1 reporter (Supplemental Fig. S9A-H’). We conclude that, also in the case of Antp auto-regulation, short linker isoforms function as more potent repressors, whereas long linker ones operate as more potent activators.

Since the Antp P1 promoter undergoes a swich from preferential auto-activation to auto-repression, and short and long linker Antp isoforms function as preferential auto-repressors and auto-activators, respectively, it appeared possible that the switch in Antp regulation is executed at the level of transcript variant abundance of these isoforms. Therefore, we next quantified the relative abundance of long and short linker transcript variants in the embryo, second and third instar discs (Fig. 4D,H). The data showed that the abundance of the long linker variant decreased, whereas the abundance of the short linker variant increased over time in development, in line with previous observations (Stroeher et al., 1988). Thus, as hypothesized, this finding suggested that relative transcript variant abundance may underlie the switch between auto-activation and auto-repression (without excluding additional mechanisms, such as changes in the chromatin modifications between early and later disc development, or the participation of different cofactors).

Relative changes in Antp transcript variant abundance (Fig. 4D,H), differential efficiency of their encoded isoforms to repress or activate the *Antp* gene (Fig. 4B-D,F-H), the developmental switch of *Antp* from auto-activation to repression (Fig. 3) and the different mobility of Antp between second and third instar imaginal discs (Fig. 3E) all pointed towards the hypothesis that the two isoforms have different modes of interaction with chromatin. To investigate this, we expressed the two isoforms from the *69B* enhancer in third instar wing and antennal discs. This results in Antp concentrations close to (if not below) endogenous levels (Supplemental Fig. S4A-J). FCS measurements revealed that the short linker isoform displayed longer characteristic decay times and a higher fraction of DNA-bound molecules, suggesting stronger and more pronounced binding to chromatin than its long linker counterpart (Fig. 4D,H and Supplemental Fig. S10A,B). With chromatin (and therefore Antp binding sites configuration), as well as the presence of cofactor proteins, being identical between the two instances (short and long linker isoforms examined in third instar wing and antennal imaginal discs of the same age), we were able to directly compare the apparent equilibrium dissociation constants for the two isoforms (Supplement 3). We found that the affinity of binding to chromatin (*K****_d_~@***) of the repressing short linker isoform is at least 2.3 times higher compared to the activating long linker isoform 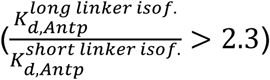 (Fig. 4D,H and Supplemental Fig. S10C-D’). Collectively, these experiments support the notion that differences in Antp regulation during disc development can be largely attributed to differences in the affinity of the investigated Antp isoforms.

Taken together, the switch of Antp from an auto-activating to an auto-repressing state and the alteration of its DNA-binding behavior during disc development can be largely explained by a temporal developmental regulation of the relative concentrations of preferentially auto-activating and auto-repressing Antp protein isoforms, which themselves display distinct properties in their modes of interaction with chromatin (Fig. 4E, I).

### Robustness of Antp auto-regulation

The mechanism of developmental Antp auto-regulation offered a possible explanation for the observed increase in Antp concentration from second to third instar discs, as well as the suppression of variability. What remained an open question is the functional significance of suppression of Antp variability in development. In order to test this in development, we require the means to manipulate variability, yet such manipulation is currently hard to achieve at the endogenous locus. However, since average concentration and variability are interdependent, we were able to use an ectopic expression system to progressively dampen Antp variability by manipulating its concentration. To this end, we expressed SynthAntp ectopically in the antennal disc, devoid of endogenous Antp expression, and monitored the extent (strength) of homeotic transformations induced by different Gal4 drivers corresponding to different SynthAntp concentrations (as measured by FCS previously in Supplemental Fig. S4A-D). In this case, expression of SynthAntp is controlled by the Gal4 driver, independently of the Antp locus, therefore the phenotypic output does not depend on Antp auto-regulation. We observed that partial transformations of antennae to tarsi could be obtained with drivers expressing Antp at close to endogenous concentration (*ptc-*Gal4, *Dll-*Gal4 (MD713) and *69B*-Gal4 drivers, Fig. 5B-D and Supplemental Fig. S4B-D). This indicates that Antp is able to repress the antennal and launch the leg developmental program in the antennal disc at endogenous concentrations, although not in a robust manner across the tissue. As expected, the three weak transformation phenotypes, elicited by *ptc-, Dll* (MD713)- and *69B*-Gal4 (Fig. 5B-D) were accompanied by high variability of SynthAntp concentration in developing discs (Fig. 5E,F). In contrast, strong expression of SynthAntp from the *Dll-*Gal4 (MD23) enhancer resulted in robust homeotic transformation to a complete tarsus (Fig. 5A) and was accompanied by low cell to cell variability (Fig. 5F). This condition resembled most closely the endogenous Antp variability in the leg disc (*CV^2^* = 0.103). Importantly, endogenous Antp and Antp overexpressed by any of the Gal4 drivers showed indistinguishable chromatin-binding behavior by FCS (Supplemental Fig. S4F and Supplemental Fig. S5A,B).

**Figure 5:**
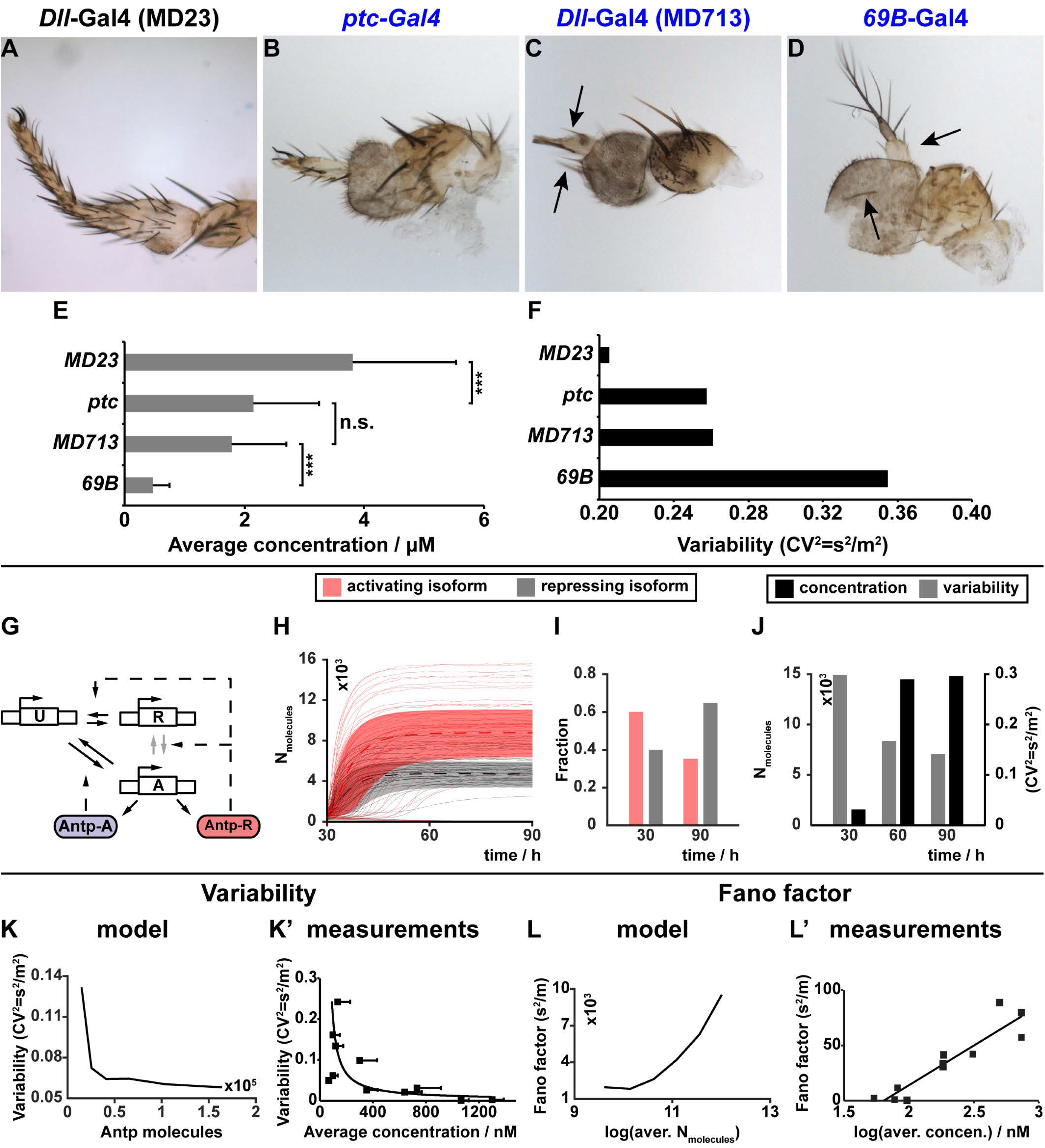
Concentrations resulting in low variability are required for Antp homeotic function. (A-D) Transformations of the distal antenna into a tarsus in adult flies, caused by *SynthAntp-eGFP* overexpression in antennal discs (<u>Supplemental Figure S4A-D</u>). Black arrows point to weaker transformations (ectopic leg bristles). (E-F) Measurements of SynthAntp concentration and cell-to-cell variability of antennal discs (<u>Supplemental Figure S4A-D</u>) in the corresponding antennal discs (A-D). (G) Dynamic Antp promoter (see text for details). (H) Trajectories of individual simulations. (I) Distribution of Antp isoforms, predicted by the model. (J) Concentration and variability predicted by the model. (K-L’) Model predictions (K and L) and experimental data validation (K’ and L’) of variability (K) and protein Fano factor (L) as a function of Antp concentration.

These results suggest that robust Antp homeotic function can be achieved at concentrations that are accompanied by low variability.

In order to further substantiate the qualitative model of Antp auto-regulation suggested by our experimental findings and to examine its impact on protein variability, we developed a simple mathematical model of stochastic *Antp* expression.

This model tests whether the identified interplay between positive and negative auto-regulation of Antp through distinct isoforms is sufficient to explain the increase in protein concentration and decrease in nucleus-to-nucleus variability from early to late stages. The model consists of a dynamic promoter, which drives transcription of *Antp* followed by a splicing step, leading to the expression of either the auto-repressing or the auto-activating isoform of Antp. In line with our finding that the repressing isoform has higher concentration at later stages, we assumed that splicing is more likely to generate this isoform than the activating isoform. The initial imbalance of Antp towards the activating isoform (Fig. 4D,H) is modeled through appropriate initial concentrations of each isoform.

Since Antp copy numbers per nucleus are in the thousands at both early and late stages, intrinsic noise of gene expression is likely to explain only a certain portion of the overall variability in Antp concentrations (Elowitz et al., 2002; Taniguchi et al., 2010). The remaining part (termed extrinsic variability) is due to cell-to-cell differences in certain factors affecting gene expression such as the ribosomal or ATP abundances. To check whether extrinsic variability significantly affects Antp expression, we expressed nuclear mRFP1 constitutively, alongside with endogenous Antp and measured the abundances of green-labeled Antp and mRFP1 (Supplemental Fig. S11). Since extrinsic factors are expected to affect both genes in a similar way, one should observe a correlation between the concentration of nuclear mRFP1 and Antp-eGFP. Our data showed a statistically significant correlation between mRFP1 and Antp (Supplemental Fig. S11C, *r =* 0.524 and *p =* 9.77 · 10^_5^). Correspondingly, we accounted for extrinsic variability also in our model by allowing gene expression rates to randomly vary between cells (Zechner et al., 2012).

The promoter itself is modeled as a Markov chain with three distinct transcriptional states. In the absence of Antp, the promoter is inactive and transcription cannot take place (state “U” in Fig. 5G). From there, the promoter can switch into a highly expressing state “A” at a rate that is assumed to be proportional to the concentration of the long-linker, auto-activating isoform (Antp-A, Fig. 5G). This resembles the positive auto-regulatory function of Antp. Conversely, the promoter can be repressed by recruitment of the short-linker, auto-repressing isoform, corresponding to state “R” in the model (Antp-R, Fig. 5G). Since the auto-repressing isoform of Antp can also activate the promoter, albeit significantly weaker than the auto-activating isoform, and vice versa, we allow the promoter to switch between states “A” and “R”.

While this promoter model resembles the dual-feedback structure of *Antp* locus inferred from experiments, it is unclear whether the two isoforms compete for the same binding sites on the P1 promoter or if auto-repression can take place regardless of whether an activating isoform is already bound to the promoter. In the former case, an increase in concentration of repressing Antp species enhances the probability to reach state “R” only if the promoter is in state “U” (Fig. 5G). In the latter case, the rate of switching between “A” and “R” also depends on the concentration of repressing isoforms of Antp (Fig. 5G, compared to Supplemental Fig. S12A). We analyzed both model variants by forward simulation and found that both of them can explain the switch-like increase in average Antp concentration between early and late stages (Fig. 5J, compared to Supplemental Fig.S12D), as well as the relative fraction of repressing and activating isoforms (Fig. 5I, compared to Supplemental Fig. S12C). However, only the non-competitive binding model (Fig. 5G) can explain the substantial reduction of total Antp variability between early and late stages (Fig. 5J), whereas in the competitive model variability is not reduced (Supplemental Fig. S12D). Simulation trajectories of individual nuclei indicated an initial increase and a subsequent stabilization of concentration, whereas in the competitive model, or in the absence of the negative feedback, this is not achieved (Fig. 5H, compared to Supplemental Fig. S12B,F). Additionally, we established that the negative feedback is required for suppression of variability (Supplemental Fig. S12E,H), since without this, no suppression of variability is conferred (Supplemental Fig. S12H). Thus, the model suggested that auto-repression is required and it is possible also if an activating isoform of Antp is already bound to the P1 promoter. Correspondingly, we use the non-competitive promoter model for further analyses.

To further validate this model, we analyzed how Antp variability scales with average concentrations and compared it to our experimental measurements. To generate different average concentrations, we varied the gene expression rates over three orders of magnitude and calculated the corresponding variability. The model predicted a decrease in variability as a function of total Antp concentration and an increase in the Fano factor. These findings are in good agreement with the experimental data (compare Fig. 5K to K’ and L to L’).

We next analyzed the model behavior under different genetic perturbations. Increase of Antp concentration by overexpressing SynthAntp transgenes (bearing either a long or a short linker isoform) from the Antp P1 promoter (Antp P1-Gal4>UAS-SynthAntp-eGFP long or short linker) resulted in 100% embryonic lethality, rendering the analysis of concentration and variability in imaginal discs impossible. This indicated that indiscriminate increase of the dosage of either Antp variant from early embryonic development onwards cannot be tolerated or buffered by the auto-regulatory circuit.

However, overexpression from a *Dll* enhancer [*Dll-*Gal4 (MD23)] in the leg discs or in the notum (*MS243*-Gal4), which overlaps with the endogenous Antp expression pattern only during first instar disc development (Emerald and Cohen, 2004), resulted in normal adult leg and notum structures. Flies overexpressing either the SynthAntp auto-activating or the auto-repressing isoform in distal appendages (Fig. 6A,B) or the notum (Supplemental Fig. S13A) displayed the wild type morphology, indicative of normal Antp function, regardless of which isoform (activating or repressing) was overexpressed. To understand this observation, we measured by FCS the concentration and variability of the total Antp protein (endogenous Antp-eGFP and overexpressed SynthAntp-eGFP) in proximal regions of the leg disc at second and third instar stages (Fig. 6C,C’). We found that, while the concentration remained high at early and later stages due to the overexpression, variability was reduced to endogenous levels at late stages. Also, the reduction of Antp variability does not seem to depend on its concentration alone, because for high concentrations at both early and late stages, variability is high only in the early stage and reduced in the late stage.

**Figure 6:**
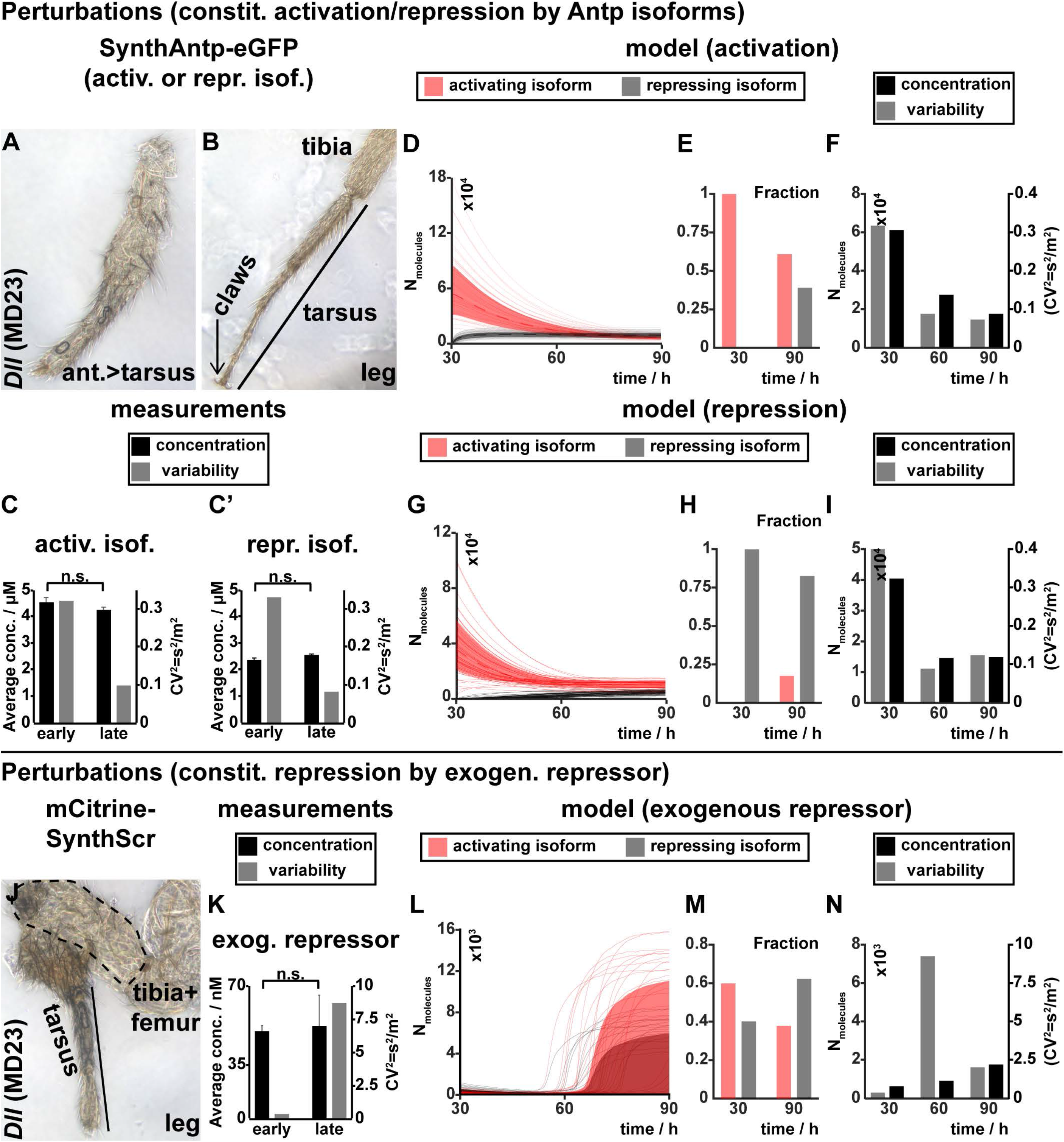
Response of *Antp* to genetic perturbations. (A-B) Overexpression of SynthAntp-eGFP long or short linker isoform result in tarsal transformations of the antenna (A), but normal leg development (B). (C-C’) Antp concentration and variability, measured by FCS, in leg discs of second and third instar larvae upon SynthAntp-eGFP long or short linker isoform expression. (D-I) Model response upon overexpression of Antp activating or repressing isoforms (similar to Fig. 5H-J). (J) Overexpression of an exogenous repressor (Scr) results in abnormal leg development Scr. (K-N) Similar to (C-I) (see also Supplemental Figure S13E-I’). (K) Antp concentration and variability, measured by FCS in the proximal leg disc of second (early) and third (late) instar larvae upon overexpression of mCherry-SynthScr.

Taken together, the phenotypic analysis and FCS measurements indicate that Antp auto-regulation is able to reduce variability, even at high levels of expression of either isoform, ensuring proper leg development.

The experimental data were corroborated by the model, which predicted that more than three-fold overexpression of either auto-activating or auto-repressing Antp isoforms will nevertheless equilibrate to normal expression levels at later stages (Fig. 6D,F,G,I). Specifically, we have measured by FCS roughly 15400 molecules in the wild type leg disc, and the model is in good quantitative agreement with this measurement upon overexpression of the activating or repressing isoform. In addition, there is no negative effect on the noise suppressing property of the circuit (Fig. 6F, I). Thus, both the model and experimental data indicate that transient high levels of either isoform early during disc development can be tolerated and that the concentration and cell-to-cell variability of the endogenous Antp protein is restored at later stages.

In contrast, overexpression of an exogenous repressor, such as Sex combs reduced (Scr), which can repress Antp at the transcriptional level, but can neither activate it nor activate its own transcription (Supplemental Fig. S13E-J’), resulted in abnormal leg (Fig. 6J) and notum (Supplemental Fig. S13B) development. These flies died as pharate adults with malformed legs that did not allow them to hatch from the pupal case, in line with Antp being required for proper leg development in all ventral thoracic discs (legs). FCS measurements in the corresponding proximal leg disc cell nuclei of second and third instar leg discs overexpressing mCherry-SynthScr revealed a pronounced reduction in Antp concentration and a remarkable increase in variability (Fig. 6K). In line with these observations, the model predicted a similar block of transcription and correspondingly severe effects on Antp dynamics (Fig. 6L-N). In both the measurements and the model prediction, the high increase in variability was triggered by the fact that a majority of the cells do not manage to switch into the highly-expressing state since too few long-linker Antp molecules are present to establish the positive auto-regulation. Since splicing favors the short-linker isoforms at later stages, these cells never “recover” from Scr repression after restriction of the Antp overexpression domain to proximal regions of the leg disc (Fig. 6L).

Taken together, the minimal model of Antp auto-regulatory genetic circuit is able to explain the experimentally observed differences in Antp concentration and cell-to-cell variability at early and late developmental stages.

## Discussion

In this work, we have characterized the endogenous molecular numbers (concentration) and cell-to-cell variability in concentration of 14 TFs in *Drosophila* imaginal discs by FCS. We have identified Antp as a TF displaying low variability among cells. We used genetics, FCS and mathematical modeling to quantitatively characterize Antp behavior in live imaginal discs and identified a kinetic mechanism responsible for the suppression of variability in the third instar discs compared to earlier developmental stages. We found that *Antp* can auto-regulate its expression levels during the course of development, starting from a preferentially auto-activating state early in development and transitioning to a preferentially auto-repressing state later. The early state is characterized by lower average Antp concentrations and high variability, whereas the opposite is true for the later repressing state. Without excluding other mechanisms, such as chromatin configuration, accessibility of Hox binding sites to Antp, the differential abundance of cofactors among developmental stages, or different modes of interactions with different Antp isoforms, we have shown that differential expression of Antp isoforms is one contributing mechanism for the observed regulatory switch. These isoforms have preferentially activating or repressing activities on the Antp promoter, bind chromatin with different affinities and are themselves expressed in different relative amounts during development. A loss-of-function analysis of the isoforms *in vivo* will be required to provide a definitive answer on the relative contribution of the Antp isoform-mediated auto-regulatory circuit towards observed suppression of variability. CRISPR/Cas9-mediated genome manipulation, in principle, allows the generation of Antp loci that express only one or the other isoform. However, it is not clear whether these flies can reach the larval developmental stages, given the Antp embryonic functions and, in fact, strong biases towards only the activating or repressing isoform introduced by Antp-Gal4-mediated expression of either Antp isoform resulted in embryonic lethality. In the absence of such direct evidence, we turned to mathematical modelling and derived, based on our experimental data, a simple kinetic model of *Antp* auto-regulation that confirmed the plausibility of the proposed mechanism. In addition, the model generated predictions that could be verified by introducing genetic perturbations.

Negative auto-regulation has been identified as a frequently deployed mechanism for the reduction of noise (cell-to-cell variability) and the increase of regulatory robustness in various systems (Becskei and Serrano, 2000; Dublanche et al., 2006; Gronlund et al., 2013; Nevozhay et al., 2009; Shimoga et al., 2013; Thattai and van Oudenaarden, 2001). Auto-repression has been described for the Hox gene *Ultrabithorax* (*Ubx*) in haltere specification and as a mechanism of controlling Ubx levels against genetic variation (Crickmore et al., 2009; Garaulet et al., 2008), as well as in *Ubx* promoter regulation in Drosophila S2 cells (Krasnow et al., 1989). In contrast, an auto-activating mechanism is responsible for the maintenance of *Deformed* expression in the embryo (Kuziora and McGinnis, 1988). Moreover, global auto-regulation of *Hox* gene complexes has been shown to be in effect also in mammalian limb development (Sheth et al., 2014). These experiments point to evolutionarily conserved mechanisms for establishing (auto-activation) or limiting (auto-repression) Hox TF levels and variability in different developmental contexts.

Our data suggest that the developmental switch from auto-activation to auto-repression is, at least in part, mediated by molecularly distinct Antp linker isoforms. Differences in affinities of different Hox TF isoforms, based on their linker between the YPWM motif and the homeodomain, have also been identified for the Hox TF Ubx. Interestingly, its linker is also subject to alternative splicing at the RNA level (Reed et al., 2010). In a similar way to Antp, the long linker Ubx isoform displays four to five fold lower affinity of DNA binding, as compared to short linker isoforms, and the two isoforms are not functionally interchangeable in *in vivo* assays. Finally, the Ubx linker also affects the strength of its interaction with the Hox cofactor Extradenticle (Exd), underscoring the functional importance of linker length in Hox TF function (Saadaoui et al., 2011). Thus, protein isoform control might represent a common regulatory mechanism of Hox-mediated transcriptional regulation.

Mathematical modeling predicts that the Antp auto-regulatory circuit is robust with respect to initial conditions and extrinsic noise by being able to suppress cell-to-cell concentration variability even at high concentrations of any of the two Antp isoforms (auto-repressing or auto-activating). This “buffering” capacity on cell-to-cell variability is reflected in the ability of flies to tolerate more than 3-fold overexpression of Antp without dramatic changes in endogenous Antp levels or generation of abnormal phenotypes. Therefore, two different isoforms produced from the same gene with opposing roles in transcriptional regulation and different auto-regulatory binding sites on the gene’s promoter seem to suffice to create a robust gene expression circuit that is able to “buffer” perturbations of the starting conditions.

So far, we have only been able to indiscriminately increase or decrease Antp concentration at the tissue level and record the phenotypic outcome of these boundary states. It will be interesting to test whether controlled perturbations of TF variability at the tissue level that render TF concentration patterns less or more noisy among neighboring cells, lead to abnormal phenotypes. The technology to selectively manipulate expression variability of specific TF in a developing tissue is yet to be established.

## Materials and Methods

### Fly stocks used

The Antp-eGFP MiMIC line has been a kind gift from Hugo J. Bellen (Bloomington stock 59790). The *atonal* (VDRC ID 318959), *brinker* (VDRC ID 318246), *spalt major* (VDRC ID 318068), *yorkie* (VDRC ID 318237), *senseless* (VDRC ID 318017) and *Sex combs reduced* (VDRC ID 318441) fosmid lines are available from the Vienna Drosophila Resource Center (VDRC) and have been generated recently in our laboratory (Sarov et al., 2016). The *fork head* (stock 43951), *grainy head* (stock 42272), *Abdominal B* (stock 38625), *eyeless,* (stock 42271), *spineless* (transcript variant A, stock 42289), and *grain* (stock 58483) tagged BACs were generated by Rebecca Spokony and Kevin P. White and are available at the Bloomington Stock Center. For the *scalloped* gene, a GFP-trap line was used (Buszczak et al., 2007), a kind gift from Allan C. Spradling laboratory (line CA07575), with which genome-wide chromatin immunoprecipitation experiments have been performed (Slattery et al., 2013). For the *spineless* gene, Bloomington stock 42676, which tags isoforms C and D of the Spineless protein has been also tried in fluorescence imaging and FCS experiments, but did not yield detectable fluorescence in the antennal disc, rendering it inappropriate to be used in our analysis. Therefore, we resided to stock 42289, which tags the A isoform of the protein. For the *eyeless* gene, the FlyFos015860(pRedFlp-Hgr)(ey13630::2XTY1 -SGFP-V5-preTEV-BLRP-3XFLAG)dFRT line (VDRC ID 318018) has been tried also in fluorescence imaging and FCS experiments, but did not yield detectable fluorescence in the eye disc for it to be used in our analysis. The *act5C*-FRT-yellow-FRT-Gal4 (Ay-Gal4) line used for clonal overexpression or *RNAi* knockdown has been described (Ito et al., 1997). The UAS-Antp lines (synthetic and full-length), as well as UAS-SynthScr constructs have been previously described (Papadopoulos et al., 2011; Papadopoulos et al., 2010). The *Dll-*Gal4 (MD23) line has been a kind gift of Ginés Morata (Calleja et al., 1996). *69B*-Gal4 and *ptc-*Gal4 have been obtained from the Bloomington Stock Center. The *Antp* P1-*lacZ* and P2-*lacZ* have been previously described (Engstrom et al., 1992; Zink et al., 1991). The P1 reporter construct spans the region between 9.4 kb upstream of the P1 promoter transcription initiation site and 7.8 kb downstream into the first intron, including the first exon sequences and thus comprising 17.2 kb of Antp regulatory sequences (pAPT 1.8). The line used has been an insertion of the pAPT 1.8 vector bearing the P1 promoter regulatory sequences upstream of an *actin-lacZ* cytoplasmic reporter and has been inserted in cytogenetic location 99F on the right chromosomal arm of chromosome 3. The *Antp-RNAi* line has been from VDRC, line KK101774. UAS-eGFP stock was a kind gift of Konrad Basler. We are indebted to Sebastian Dunst for generating the *ubi*-FRT-mCherry(stop)-FRT-Gal4(VK37)/CyO line, which drives clonal overexpression upon flippase excision, while simultaneously marking cells by the loss of mCherry. For red-color labeling of clones the *act5C*-FRT-CD2-FRT-Gal4, UAS-mRFP1(NLS)/TM3 stock 30558 from the Bloomington Stock Center has been used. For marking the ectopic expression domain of untagged Antp proteins the UAS-mRFP1(NLS)/TM3 stock 31417 from the Bloomington Stock Center has been used. The *MS243*-Gal4; UAS-GFP/CyO line was a kind gift from the laboratory of Ernesto Sánchez-Herrero.

### Fly genotypes corresponding to fluorescence images

Supplemental Fig. S1A: FlyFos018487(pRedFlp-Hgr)(ato37785::2XTY1-SGFP-V5-preTEV-BLRP-3XFLAG)dFRT

Supplemental Fig. S1B: FlyFos024884(pRedFlp-Hgr)(brk25146::2XTY1-SGFP-V5-preTEV-BLRP-3XFLAG)dFRT

Supplemental Fig. S1C: FlyFos030836(pRedFlp-Hgr)(salm30926::2XTY1-SGFP-V5-preTEV-BLRP-3XFLAG)dFRT

Supplemental Fig. S1: FlyFos029681(pRedFlp-Hgr)(yki19975::2XTY1-SGFP-V5-preTEV-BLRP-3XFLAG)dFRT

Supplemental Fig. S1E: w^1118^; PBac(fkh-GFP.FPTB)VK00037/SM5 Supplemental Fig. S1F: *sd*-eGFP (FlyTrap, homozygous)

Supplemental Fig. S1G: w^1118^; PBac(grh-GFP.FPTB)VK00033

Supplemental Fig. S1H: FlyFos018974(pRedFlp-Hgr)(Scr19370::2XTY1-SGFP-V5-preTEV-BLRP-3XFLAG)dFRT

Supplemental Fig. S1I: FlyFos015942(pRedFlp-Hgr)(sens31022::2XTY1-SGFP-V5-preTEV-BLRP-3XFLAG)dFRT

Supplemental Fig. S1J,K: Antp-eGFP (MiMIC) homozygous (line MI02272, converted to an artificial exon)

Supplemental Fig. S1L: w^1118^; PBac(Abd-B-EGFP.S)VK00037/SM5

Supplemental Fig. S1M: w^1118^; PBac(ey-GFP.FPTB)VK00033

Supplemental Fig. S1N: w^1118^; PBac(ss-GFP.A.FPTB)VK00037

Supplemental Fig. S1O,P: w^1118^; PBac(grn-GFP.FPTB)VK00037

Fig. 2B,B’: *hs*-flp/+; *act5C*-FRT-yellow-FRT-Gal4/+; UAS-SynthAntp long linker-eGFP/+

Fig. 2C,C’: *hs*-flp/+; *act5C*-FRT-yellow-FRT-Gal4, UAS-eGFP/+; UAS-Antp long linker (full-length, untagged)/+

Fig. 2F,F’: *Dll-*Gal4 (MD23)/+; UAS-SynthAntp-eGFP/*Antp* P1-*lacZ* Supplemental Fig. S3A,A’: *Antp* P1-*lacZ*/TM3

Supplemental Fig. S3B,B’: *Antp* P2-*lacZ*/GyO

Supplemental Fig. S3C,C’: wild type

Supplemental Fig. S3D,D’: *hs*-flp; *act5C*-FRT-yellow-FRT-Gal4, UAS-eGFP

Supplemental Fig. S3E,E’: *hs*-flp/+; *act5C*-FRT-yellow-FRT-Gal4, UAS-eGFP/+; *Antp* P1-*lacZ*/+

Supplemental Fig. S3F,F’: *hs*-flp/+; *act5C*-FRT-yellow-FRT-Gal4, UAS-eGFP/+; UAS-Antp long linker (full-length, untagged)/*Antp* P1-*lacZ*

Supplemental Fig. S3G,G’: *Dll-*Gal4 (MD23)/+; UAS-Antp long linker (full-length, untagged), UAS-mRFP1(NLS)/ *Antp* P1-*lacZ*

Supplemental Fig. S3H,H’: *Dll-*Gal4 (MD23)/+; UAS-mRFP1(NLS)/ *Antp* P1-*lacZ*

Supplemental Fig. S4A: *Dll-*Gal4 (MD23)/+; UAS-SynthAntp long linker-eGFP/+

Supplemental Fig. S4B: *ptc-*Gal4/+; UAS-SynthAntp long linker-eGFP/+

Supplemental Fig. S4C: *Dll-*Gal4 (MD713)/+; UAS-SynthAntp long linker-eGFP/+

Supplemental Fig. S4D,G,H,K: *69B*-Gal4/UAS-SynthAntp long linker-eGFP

Supplemental Fig. S4I,J,L: *69B*-Gal4/UAS-eGFP

Fig. 3B,B’,G,G’: *hs*-flp/+; *ubi*-FRT-mChery-FRT-Gal4/+; Antp-eGFP (MiMIC)/UAS-Antp long linker (full-length, untagged)

Fig. 3C,C’: *hs*-flp/+; UAS-Antp*^RNAi^*/+; *Antp* P1-*lacZ*/*act5C*-FRT-CD2-FRT-Gal4, UAS-mRFP1(NLS)

Fig. 3H,H’: *hs*-flp/+; UAS-Antp*^RNAi^*/*act5C*-FRT-yellow-FRT-Gal4, UAS-eGFP; *Antp* P1- *lacZ*/*+*

Supplemental Fig. S6B: *Antp* P1-*lacZ*/TM6B

Supplemental Fig. S7A,A’: *hs*-flp/+; *ubi*-FRT-mChery-FRT-Gal4/+; Antp-eGFP (MiMiG)/UAS-Antp long linker (full-length, untagged)

Supplemental Fig. S7B-C’: *hs*-flp/+; *ubi*-FRT-mChery-FRT-Gal4/+; Antp-eGFP (MiMIC)/+

Supplemental Fig. S7D,D’: *hs*-flp/+; *act5C*-FRT-yellow-FRT-Gal4, UAS-eGFP/+; *Antp* P1-*lacZ*/UAS-Antp long linker (full-length, untagged)

Supplemental Fig. SE,E’: *hs*-flp/+; *act5C*-FRT-yellow-FRT-Gal4/+; UAS-SynthAntp long linker-eGFP/+

Supplemental Fig. S7F,F’: *hs*-flp/+; *act5C*-FRT-yellow-FRT-Gal4, UAS-eGFP/+

Supplemental Fig. S7G,G’: *hs*-flp/+; *act5C*-FRT-yellow-FRT-Gal4, UAS-eGFP/+; *Antp* P1-*lacZ*/+

Supplemental Fig. S7H,H’: *hs*-flp/+; UAS-Antp*^RNAi^*/+; Antp-eGFP (MiMIC)/*act5C*-FRT-CD2-FRT-Gal4, UAS-mRFP1(NLS)

Fig. 4B,B’: *ptc-*Gal4/+; UAS-SynthAntp long linker-eGFP/+

Fig. 4C,C’: *Dll-*Gal4 (MD23)/+; UAS-SynthAntp long linker-eGFP/*Antp* P1-*lacZ*

Fig. 4F,F’: *ptc-*Gal4/+; UAS-SynthAntp long linker-eGFP/+

Fig. 4G,G’: *Dll-*Gal4 (MD23)/+; UAS-SynthAntp short linker-eGFP/*Antp* P1-*lacZ*

Supplemental Fig. S9A,A’: *hs*-flp/+; *act5C*-FRT-yellow-FRT-Gal4/+; UAS-SynthAntp short linker-eGFP/+

Supplemental Fig. S9B,B’,G,G’: *hs*-flp/+; *act5C*-FRT-yellow-FRT-Gal4/+; UAS-SynthAntp short linker-eGFP/*Antp* P1-*lacZ*

Supplemental Fig. S9C,C’,H,H’: *hs*-flp/+; *act5C*-FRT-yellow-FRT-Gal4/+; UAS-Antp short linker (full-length, untagged)/*Antp* P1-*lacZ*

Supplemental Fig. S9D,D’: *hs*-flp/+; *Dll-*Gal4 (MD23)/+; UAS-Antp short linker (full-length, untagged), UAS-mRFP1(NLS)/*Antp* P1-*lacZ*

Supplemental Fig. S9E-F’: *hs*-flp/+; *ubi*-FRT-mChery-FRT-Gal4/+; Antp-eGFP (MiMIC)/UAS-Antp short linker (full-length, untagged)

Supplemental Fig. S9I,I’: *ptc-*Gal4/+; UAS-SynthAntp long linker-eGFP/*Antp* P1-*lacZ*

Supplemental Fig. S9J,J’: *ptc-*Gal4/+; UAS-SynthAntp short linker-eGFP/*Antp* P1-*lacZ*

Fig. 5A: *Dll-*Gal4 (MD23)/+; UAS-SynthAntp long linker-eGFP/+

Fig. 5B: *ptc-*Gal4/+; UAS-SynthAntp long linker-eGFP/+

Fig. 5C: *Dll-*Gal4 (MD713)/+; UAS-SynthAntp long linker-eGFP/+

Fig. 5D: *69B*-Gal4/UAS-SynthAntp long linker-eGFP

Supplemental Fig. S11A-B’: *ubi*-mRFP1(NLS)/+ or y; Antp-eGFP (MiMIC)/+

Supplemental Fig. S12B,C: *Dll-*Gal4 (MD23)/+; UAS-mCitrine-SynthScr/+

Fig. 6A,B: *Dll-*Gal4 (MD23)/+; UAS-SynthAntp long linker-eGFP/+ or *Dll-*Gal4 (MD23)/+; UAS-SynthAntp short linker-eGFP/+

Fig. 6J: *Dll-*Gal4 (MD23)/+; UAS-mCitrine-SynthScr/+

Supplemental Fig. S13A,D,D’: *MS243*-Gal4/+; UAS-SynthAntp long linker-eGFP/*Dr* or *MS243*-Gal4/+; UAS-SynthAntp short linker-eGFP/*Dr*

Supplemental Fig. S13B,F,F’: *MS243*-Gal4/+; UAS-mCitrine-SynthScr/+

Supplemental Fig. S13C,I,I’: *Dll-*Gal4 (MD23)/+; UAS-mCitrine-SynthScr/+

Supplemental Fig. S13E,E’: *ptc-*Gal4/+; UAS-SynthAntp long linker-eGFP/+

Supplemental Fig. S13F,F’: *MS243*-Gal4/+; UAS-mCitrine-SynthScr/+

Supplemental Fig. S13G,G’: *ptc-*Gal4/+; UAS-mCitrine-SynthScr/*Antp* P1*-lacZ*

Supplemental Fig. S13H,H’: *Dll-*Gal4 (MD23)/+; UAS-mCitrine-SynthScr/*Antp* P1-*lacZ*

Supplemental Fig. S13J,J’: *MS243*-Gal4/+; UAS-eGFP/+

### Preparation of second and third instar imaginal discs for FCS measurements

For FCS measurements, imaginal discs (eye-antennal, wing, leg, humeral and genital) and salivary glands were dissected from third instar wandering larvae, or wing and leg discs from second instar larvae, in Grace’s insect tissue culture medium (ThermoFisher Scientific, 11595030) and transferred to 8-well chambered coverglass (Nunc® Lab-Tek™, 155411) containing PBS just prior to imaging or FCS measurements. Floating imaginal discs or salivary glands were sunk to the bottom of the well using forceps.

### Immunostainings in larval imaginal discs

Larval imaginal discs were stained according to (Papadopoulos et al., 2010). Stainings for the endogenous Antp protein have been performed using a mouse anti-Antp antibody (Developmental Studies Hybridoma Bank, University of Iowa, anti-Antp 4C3) in a dilution of 1:250 for embryos and 1:500 for imaginal discs. eGFP, or eGFP-tagged proteins have been stained using mouse or rabbit anti-GFP antibodies from ThermoFisher Scientific in a dilution of 1:500 in imaginal discs and 1:250 in embryos. mRFP1 was stained using a Chromotek rat anti-RFP antibody. For *Antp* P1 promoter stainings in imaginal discs we used the mouse anti-β-galactosidase 40-1a antibody from Developmental Studies Hybridoma Bank, University of Iowa in a dilution of 1:50. The rabbit anti-Scr antibody was used in a dilution of 1:300 (LeMotte et al., 1989). Confocal images of antibody stainings represent predominatly Z-projections and Zeiss LSM510, Zeiss LSM700 or Zeiss LSM880 Airyscan confocal laser scanning microscopy systems with an inverted stand Axio Observer microscope were used for imaging. Image processing and quantifications have been performed in Fiji (Schindelin et al., 2012). For optimal spectral separation, secondary antibodies coupled to Alexa405, Alexa488, Alexa594 and Cy5 (ThermoFischer Scientific) were used.

### Colocalization of wild type and eGFP-tagged MiMIC Antp alleles in imaginal discs

To examine whether the pattern of the MiMIC Antp-eGFP fusion protein recapitulates the Antp wild type expression pattern in both embryo and larval imaginal discs, we performed immunostainings of heterozygous Antp-eGFP and wild type flies to visualize the embryonic (stage 13) and larval expression of *Antp* and eGFP. In this experiment, we 1) visualized the overlap between eGFP and *Antp* (the eGFP pattern reflects the protein encoded by the MiMIC allele, whereas the *Antp* pattern reflects the sum of protein produced by the MiMIC allele and the allele of the balancer chromosome) and 2) compared the eGFP expression pattern to the Antp expression pattern in wild type discs and embryos.

### Induction of early and late overexpression and *RNAi*-knockdown clones in imaginal discs

Genetic crosses with approximately 100 virgin female and 100 male flies were set up in bottles and the flies were allowed to mate for 2 days. Then, they were transferred to new bottles and embryos were collected for 6 hours at 25°C. Flies were then transferred to fresh bottles and kept until the next collection at 18°C. To asses Antp auto-activation, the collected eggs were allowed to grow at 25°C for 26 h from the midpoint of collection, when they were subjected to heat-shock by submersion to a water-bath of 38°C for 30 min and then placed back at 25°C until they reached the stage of third instar wandering larvae, when they were collected for dissection, fixation and staining with antibodies. To assess Antp auto-repression, the same procedure was followed, except that the heat-shock was performed at 60 h of development after the midpoint of embryo collection. Whenever necessary, larval genotypes were selected under a dissection stereomicroscope with green and red fluorescence filters on the basis of *deformed* (*dfd*)-YFP bearing balancer chromosomes (Le et al., 2006) and visual inspection of fluorescence in imaginal discs.

### Measurement of Antp transcript variant abundance

The linker between the Antp YPWM motif and the homeodomain contains the sequence RSQFGKCQE. Short linker isoforms encode the sequence RSQFE, whereas long linker isoforms are generated by alternative splicing of a 12 base pair sequence encoding the four amino acid sequence GKCQ into the mRNA. We initially designed primer pairs for RT-qPCR experiments to distinguish between the short and long linker mRNA variants. For the short linker variant, we used nucleotide sequences corresponding to RSQFERKR (with RKR being the first 3 amino acids of the homeodomain). For detection of the long linker variant we designed primers either corresponding to the RSQFGKCQ sequence, or GKCQERKR. We observed in control PCRs (using plasmid DNA harboring either a long or a short linker cDNA) that primers designed for the short linker variant still amplified the long linker one. Moreover, with linker sequences differing in only four amino acids, encoded by 12 base pars, primer pairs flanking the linker could also not be used, since, due to very similar sizes, both variants would be amplified in RT-qPCR experiments with almost equal efficiencies. Therefore, we used primer pairs flanking the linker region to indiscriminately amplify short and long linker variants, using non-saturating PCR (18 cycles) on total cDNA generated from total RNA. We then resolved and assessed the relative amounts of long and short linker amplicons in a second step using Fragment Analyzer (Advanced Analytical). RNA was extracted from stage 13 embryos, second instar larvae at 60 h of development, and leg or wing discs from third instar wandering larvae using the Trizol® reagent (ThermoFischer Scientific), following the manufacturer’s instructions.

Total RNA amounts were measured by NanoDrop and equal amounts were used to synthesize cDNA using High-Capacity RNA-to-cDNA™ Kit (ThermoFischer Scientific), following the manufacturer’s instructions. Total cDNA yields were measured by NanoDrop and equal amounts were used in PCR, using in-house produced Taq polymerase. 10 ng of plasmid DNA, bearing either a long or a short transcript cDNA were used as a control. PCR product abundance was analyzed both by agarose gel electrophoresis and using Fragment Analyzer (Advanced Analytical).

The quantification of the transcript variant concentration (Fig. 4 D and H) has been made considering 100% (value equal to 1 on the y axis) as the sum of long and short isoforms at each developmental stage, whereas the quantification of the relative activation and repression efficiency has been performed considering the short linker variant as having 100% repression and the long linker variant as having 100% activation (values equal to 1 on the y-axis) efficiency.

### Quantification of the relative repressing and activating efficiencies of different Antp isoforms

Quantification of the relative efficiency of Antp activating and repressing isoforms (Fig. 4D,H) were performed in Fiji (Schindelin et al., 2012) by outlining the total region of repression or activation of Antp protein or P1 reporter staining and quantification of the relative fluorescence intensity of the selected regions. From the calculated values, we have subtracted the values obtaining by outlining and calculating Antp protein or reporter beta-galactosidase staining background in the region of expression of an eGFP transgene alone (negative control). 5–7 imaginal disc images per investigated genotype were used for analysis. For the repression assay the obtained values have been normalized over the intensity of Antp protein calculated in the region of overlap between an eGFP expressing transgene and Antp (negative control). In both cases (repression and activation), the highest efficiency per transcript variant (for repression, the short linker isoform; for activation the long linker isoform) have been set to 100%.

### Fluorescence Microscopy Imaging of live imaginal discs and FCS

Fluorescence imaging and FCS measurements were performed on two uniquely modified confocal laser scanning microscopy systems, both comprised of the ConfoCor3 system (Carl Zeiss, Jena, Germany) and consisting of either an inverted microscope for transmitted light and epifluorescence (Axiovert 200 M); a VIS-laser module comprising the Ar/ArKr (458, 477, 488 and 514 nm), HeNe 543 nm and HeNe 633 nm lasers and the scanning module LSM510 META; or a Zeiss LSM780 inverted setup, comprising Diode 405 nm, Ar multiline 458, 488 and 514 nm, DPSS 561 nm and HeNe 633 nm lasers. Both instruments were modified to enable detection using silicon Avalanche Photo Detectors (SPCM-AQR-1X; PerkinElmer, USA) for imaging and FCS. Images were recorded at a 512×512 pixel resolution. C-Apochromat 40x/1.2 W UV-VIS-IR objectives were used throughout. Fluorescence intensity fluctuations were recorded in arrays of 10 consecutive measurements, each measurement lasting 10 s. Averaged curves were analyzed using the software for online data analysis or exported and fitted offline using the OriginPro 8 data analysis software (OriginLab Corporation, Northampton, MA). In either case, the nonlinear least square fitting of the autocorrelation curve was performed using the Levenberg-Marquardt algorithm. Quality of the fitting was evaluated by visual inspection and by residuals analysis. Control FCS measurements to asses the detection volume were routinely performed prior to data acquisition, using dilute solutions of known concentration of Rhodamine 6G and Alexa488 dyes. The variability between independent measurements reflects variabilitys between cells, rather than imprecision of FCS measurements. For more details on Fluorescence Microscopy Imaging and FCS, refer to Supplement 1.

In Figure 1A-H the workflow of FCS measurements is schematically represented. Live imaging of imaginal discs, expressing endogenously-tagged TFs, visualized by fluorescence microscopy and neighboring cells, expressing TFs at different levels, selected for FCS measurements (Fig. 1A-B). FCS measurements are performed by placing the focal point of the laser light into the nucleus (Fig 1C-D) and recording fluorescence intensity fluctuations (Fig. 1E), generated by the increase or decrease of the fluorescence intensity, caused by the arrival or departure of fast- and slowly-diffusing TF molecules into or out of the confocal detection volume (Fig. 1D). The recorded fluctuations are subjected to temporal autocorrelation analysis, which generates temporal autocorrelation curves (henceforth referred to as FCS curves), which by fitting with an appropriate model (Supplement 1), yield information about the absolute concentration of fluorescent molecules (F) and, after normalization to the same amplitude, their corresponding diffusion times, as well as the fraction of fast- and slowly-diffusing TF molecules (Fig. 1G). The concentration of molecules is inversely proportional to the y-axis amplitude at the origin of the FCS curve (Fig. 1F). Processes that slow down the diffusion of TF molecules, such as binding to very large molecules (e.g. chromosomal DNA), are visible by a shift of the FCS curves to longer characteristic times (Fig. 1G). Measurements in a collection of neighboring cell nuclei also allow the calculation of protein concentration variability at the live tissue level (Fig. 1H).

### Sample size, biological and technical replicates

For the measurement of TF molecular numbers and variability (Fig. 1 and Supplemental Fig. S1), 7–10 larvae of each fly strain were dissected, yielding at least 15 imaginal discs, which were used in FCS analysis. For the Fkh TF, 7 pairs of salivary glands were analyzed and for AbdB, 12 genital discs were dissected from 12 larvae. More than 50 FCS measurements were performed in patches of neighboring cells of these dissected discs, in the regions of expression indicated in Supplemental Fig. S1 by arrows. Imaginal discs from the same fly strain (expressing a given endogenously-tagged TF) were analyzed on at least 3 independent instances (FCS sessions), taking place on different days (biological replicates) and for Antp, which was further analyzed in this study, more than 20 independent FCS sessions were used. As routinely done with FCS measurements in live cells, these measurements were evaluated during acquisition and subsequent analysis and, based on their quality (high counts per molecule and second, low photobleaching), were included in the calculation of concentration and variability. In Supplemental Fig. S1Q, *n* denotes the number of FCS measurements included in the calculations.

For experiments involving immunostainings in imaginal discs to investigate the auto-regulatory behavior of Antp (Figs. 2–5 and supplements thereof, except for the temporally-resolved auto-activating and repressing study of Antp in Fig. 3, as discussed above), 14–20 male and female flies were mated in bottles and 10 larvae were selected by means of fluorescent balancers and processed downstream. Up to 20 imaginal discs were visualized by fluorescence microscopy and high resolution Z-stacks were acquired for 3–5 representative discs or disc regions of interest per experiment. All experiments were performed in triplicate, except for the temporal analysis of Antp auto-regulatory behavior in Fig. 3 (and Supplemental Figs. thereof), which was performed 6 times and the quantification of repression efficiency of short and long linker Antp isoforms in Fig. 4 (and Supplemental Figs. thereof), which was performed 5 times.

For the quantification of transcript variant abundance in Fig. 4D,H, RNA and thus cDNA was prepared from each stage 3 independent times (biological replicates) and the transcript abundance per RNA/cDNA sample was also analyzed 3 times.

For the experiments involving perturbations in Antp expression whereby the proper development of the leg and the notum have been assessed in Fig. 5, more than 100 adult flies have been analyzed and this experiment has been performed more than 10 times independently.

### Statistical significance

Fig. 3E: Statistical significance was determined using a two-tailed Student’s T-test (***, ***p < 0.001*** and *, ***p< 0.05***, namely 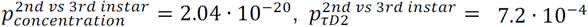 and 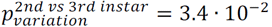).

Fig. 4D, H: Statistical significance was determined using a two-tailed Student’s T-test between measurements performed with the long linker (auto-activating) isoform (Fig. 4D) and the short linker (auto-repressing) isoform (Fig. 4H) (***, ***p <*** 0.001 and *, *p <* 0.05, namely 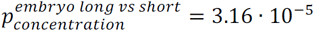, 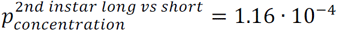,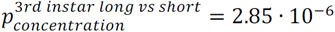, 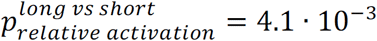, 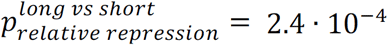 and 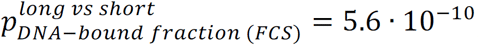).

Fig. 6C-C’: Statistical significance was determined using a two-tailed Student’s T-test (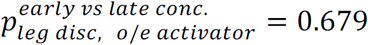 and 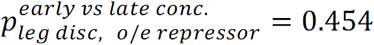).

Fig. 6K: Statistical significance was determined using a two-tailed Student’s T-test 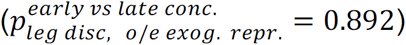.

## Acknowledgements

We are deeply saddened by the unexpected passing of Prof. Walter J. Gehring, at the very inception of this work, when the project was still in the planning and preliminary data gathering stage. Prof. Gehring was an extraordinary human being and a scientific giant, whose work will continue to educate and inspire generations to come. DKP has been supported by a long-term fellowship from the Swiss National Science Foundation (PBBSP-138700) and a long-term fellowship from the Federation of European Biochemical Societies (FEBS). VV has been supported by the Knut and Alice Wallenberg foundation and Karolinska Institute Research Funds. DKP would like to express his gratitude to PT for outstanding scientific, and uninterrupted financial, support. DKP would like to acknowledge Markus Burkhardt, Sylke Winkler, Aliona Bogdanova and the Light Microscopy facility of MPI-CBG. DKP is also grateful to Konstantinos Papadopoulos for advice on the mathematical analysis. The authors thank Wendy Bickmore, Jan Brugues and Thomas M. Schultheiss for critical comments on the manuscript.

---

**Supplemental Figure S1:**
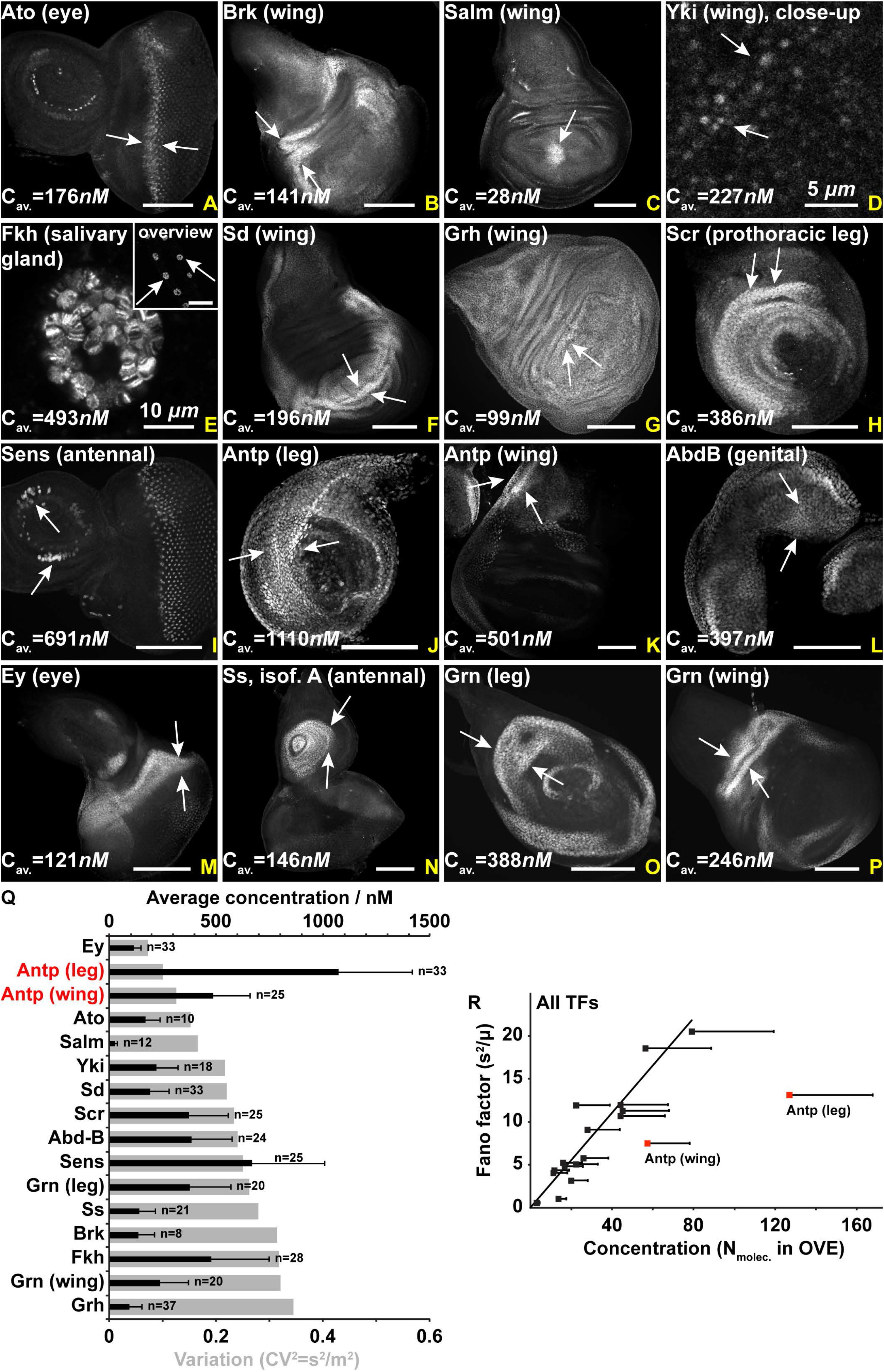
Measurement of average concentrations and nucleus-to-nucleus variability of 14 endogenously-tagged TFs in Drosophila imaginal discs by FCS. (A-P) Fluorescence imaging of TFs, showing their expression pattern in imaginal discs and the salivary gland. White arrows indicate regions where FCS measurements of endogenous intra-nuclear concentration were performed and the average concentrations are given for each TF. Images have been contrasted for visualization purposes. For the Antp and Grn TFs, both leg and wing imaginal discs have been used for measurements. Average concentrations of TFs measured in different cells span a range of two orders of magnitude, from few tens to a thousand nanomolar. Scale bars denote 100 *μm,* unless otherwise indicated. (Q) Characterization of nucleus-to-nucleus variability among neighboring cells within the same expression domain in imaginal discs of the 14 TF studied by FCS. Black bars show concentration averages (with error bars representing 1 standard deviation), whereas grey bars show the variability, i.e. the squared coefficient of variability (expressed as the variance over the squared mean, 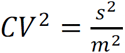). TFs have been sorted according to increasing variability. (P) Characterization of variability as a function of concentration, using the Fano factor value (expressed as variance over the mean, 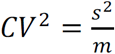). The red squares point to the Fano factor values of Antp in the wing and leg disc.

**Supplemental Figure S2:**
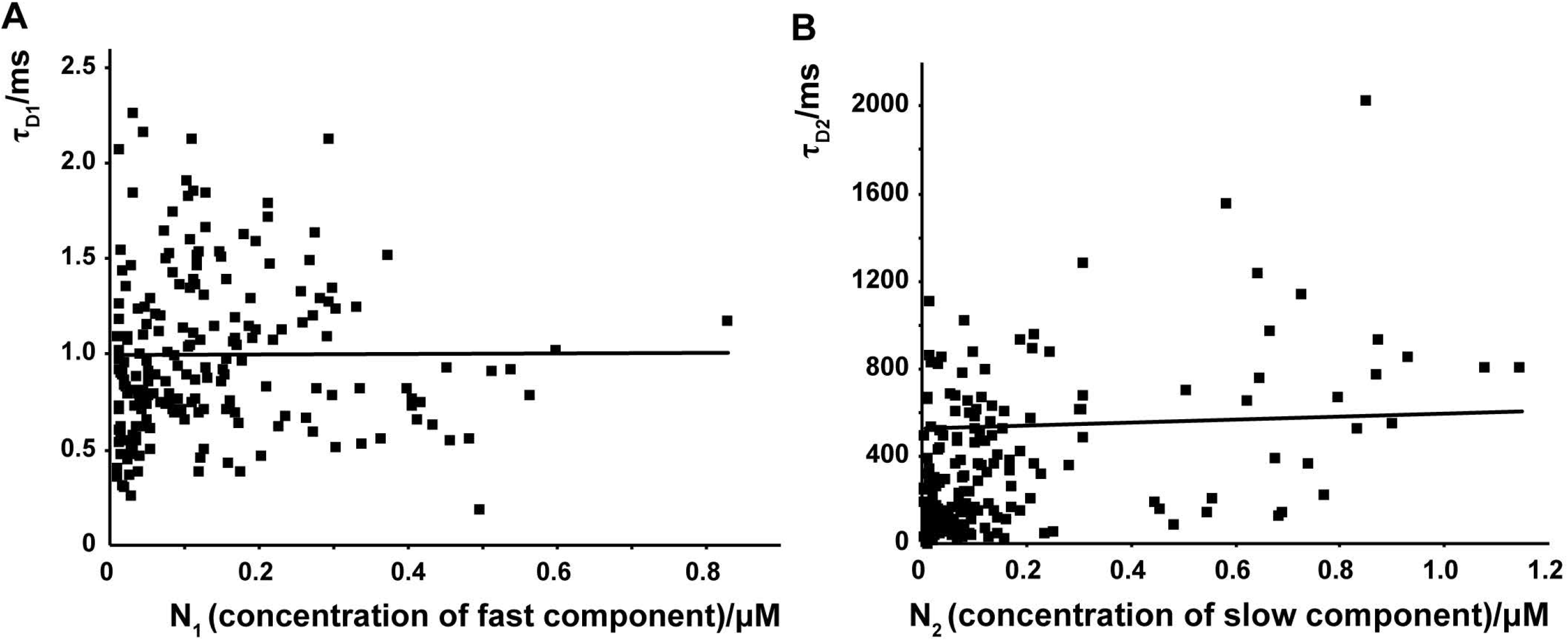
Characteristic decay times of Antp-eGFP do not change as a function of total concentration. (A-B) Characteristic decay times 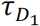 (A) and 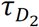 (B) do not vary with the concentration of Antp-eGFP TF molecules, as evident from 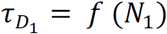 and 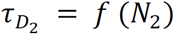, where *N*_1_ is the number of freely diffusing, *N*_2_ the number of bound Antp-eGFP TF molecules and 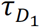, 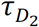 their respective diffusion times.

**Supplemental Figure S3:**
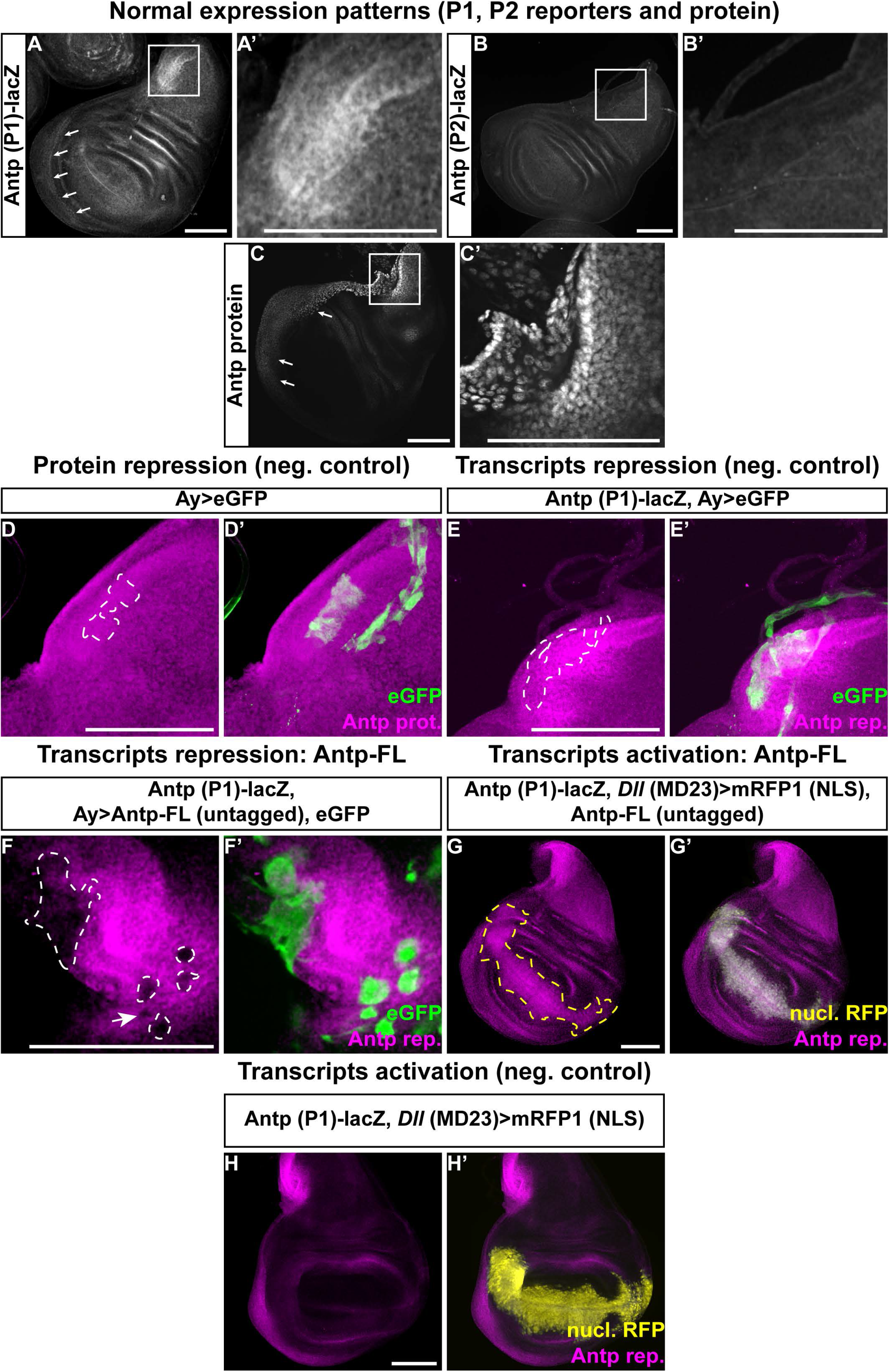
Antp is able to repress and activate itself at the transcriptional level – controls. (A-C’) Normal expression patterns of the Antp P1 (A-A’) and P2 (B-B’) transcriptional reporters and Antp protein immunohistochemistry (C-C’). Boxed areas in (A), (B) and (C) are magnified in (A’), (B’) and (C’). The Antp P1 reporter is highly expressed in the prescutum region of the notum (A’) and the peripodial cells at the base of the wing blade (giving rise to the mesopleura and pteropleura of the thorax, white arrows in (A)), which overlaps with the Antp protein pattern ((C’) and arrows in (C)). The Antp P2 promoter reporter construct exhibits very weak, if any, expression at these two domains (B-B’). (D-E’) Negative controls of Antp protein (D-D’) and P1 reporter transcription (E-E’) upon overexpression of eGFP. Dashed lines outline the regions of clonal induction in (D) and (E), where neither the Antp protein (D) nor the Antp P1 reporter (E) are repressed. (F-F’) Repression of Antp P1 reporter transcription upon clonal overexpression of the full-length untagged Antp protein (Antp-FL). The ectopic expression domain is outlined by white dashed lines in (F) and marked by the expression of eGFP (F’). (G-G’) Activation of Antp P1 reporter transcription upon ectopic expression of untagged Antp full-length (Antp-FL) with *Dll* (MD23) driver in the distal region of wing pouch. The ectopic expression domain is outlined by a yellow dashed line in (G) and is marked by the expression of nuclear mRFP1. (H-H’) Negative control of ectopic activation of Antp P1 transcription upon overexpression of nuclear mRFP1 alone by *Dll* (MD23)-Gal4. Scale bars denote 100 *μm*.

**Supplemental Figure S4:**
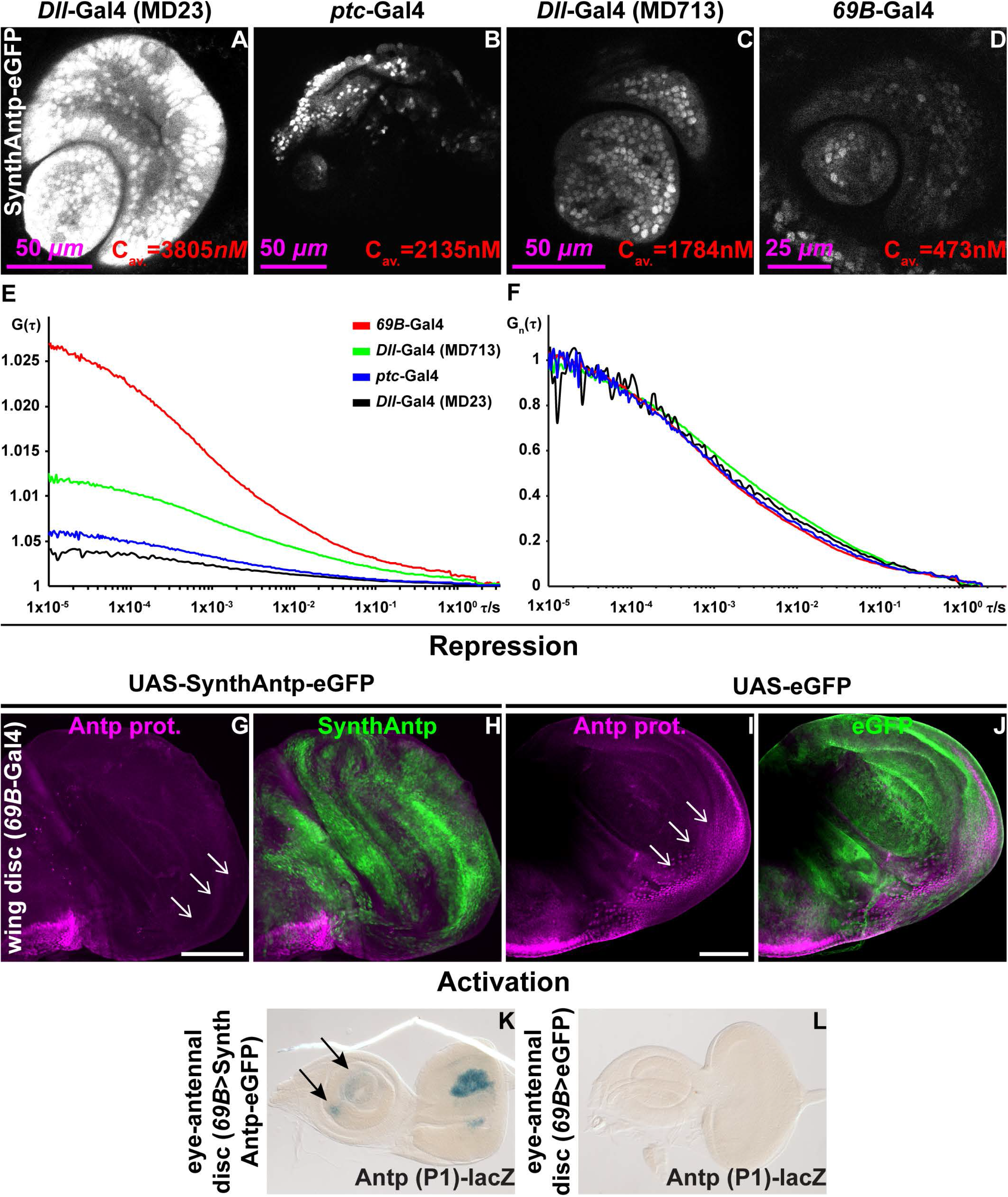
Direct correlation between Antp concentration and homeotic function – Antp auto-repression and activation occurs at endogenous concentrations. (A-D) Live imaging (one optical section) of SynthAntp-eGFP expressed in the distal antennal portion of the eye-antennal disc by different Gal4 drivers. The concentration was measured using FCS and average concentrations are indicated. An eightfold difference was observed between the strong *Dll-*Gal4 driver (MD23) (A) and weak *69B*-*Gal4* driver (D). (E) Average FCS measurements performed in nuclei overexpressing *SynthAntp-eGFP,* using different Gal4 drivers. Note that the y-axis amplitudes at the origin of the FCS curves are inversely proportional to the concentration. (F) FCS curves of measurements in (E) normalized to the same amplitude, *G_n_*(*τ*) = 1 at τ = 10 μ*s*, show major overlap, indicating indistinguishable behavior of Antp binding to chromatin across the concentration range examined (0.5 – 3.8 nM). (G-L) Antp auto-regulation occurs at endogenous concentrations. (G-H) Repression of endogenous Antp protein upon induction of *SynthAntp-eGFP* in the proximal regions of the wing disc by *69B*-*Gal4,* which results in Antp expression very similar to endogenous levels. (I-J) No repression is observed upon overexpression of *eGFP* (negative control), as indicated by white arrows in (I). White arrows in (G) and (I) point to the equivalent area in the wing disc, where Antp repression is observed. (K) X-gal stainings of the *Antp* P1 reporter show weak but detectable ectopic β-galactosidase activity in the antennal disc (black arrows). (L) Negative control stainings of eGFP induced by the *69B* enhancer show complete absence of ectopic reporter transcription. Scale bars denote 100 *μm*, unless otherwise indicated.

**Supplemental Figure S5:**
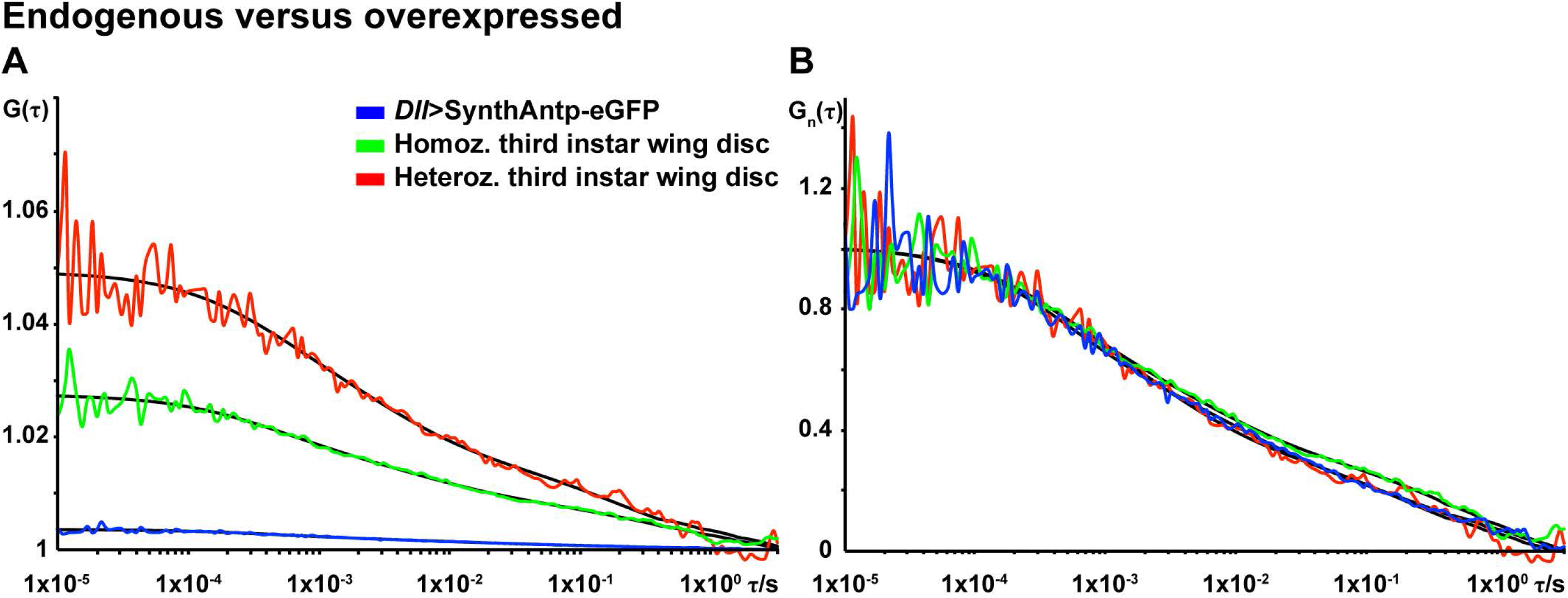
Comparison of endogenous and overexpressed Antp by FCS. (A) FCS curves of Antp-eGFP in wing disc nuclei. Concentration differences of fluorescent Antp protein are obvious among cells expressing one or two copies of Antp-eGFP (homozygous and heterozygous larvae) or overexpressing SynthAntp-eGFP from the *Dll* MD23)-Gal4 driver. (B) FCS curves shown in (A) normalized to the same amplitude, *G_n_*(*τ*) = 1 at *τ* = 10 μ*s*, show pronounced overlap between homozygous and heterozygous Antp-eGFP-expressing cells, as well as between endogenously expressed *Antp* and overexpressed *SynthAntp-eGFP,* indicating similar diffusion times and modes of interaction with chromatin. FCS curves are color-coded as outlined in panel (B).

**Supplemental Figure S6:**
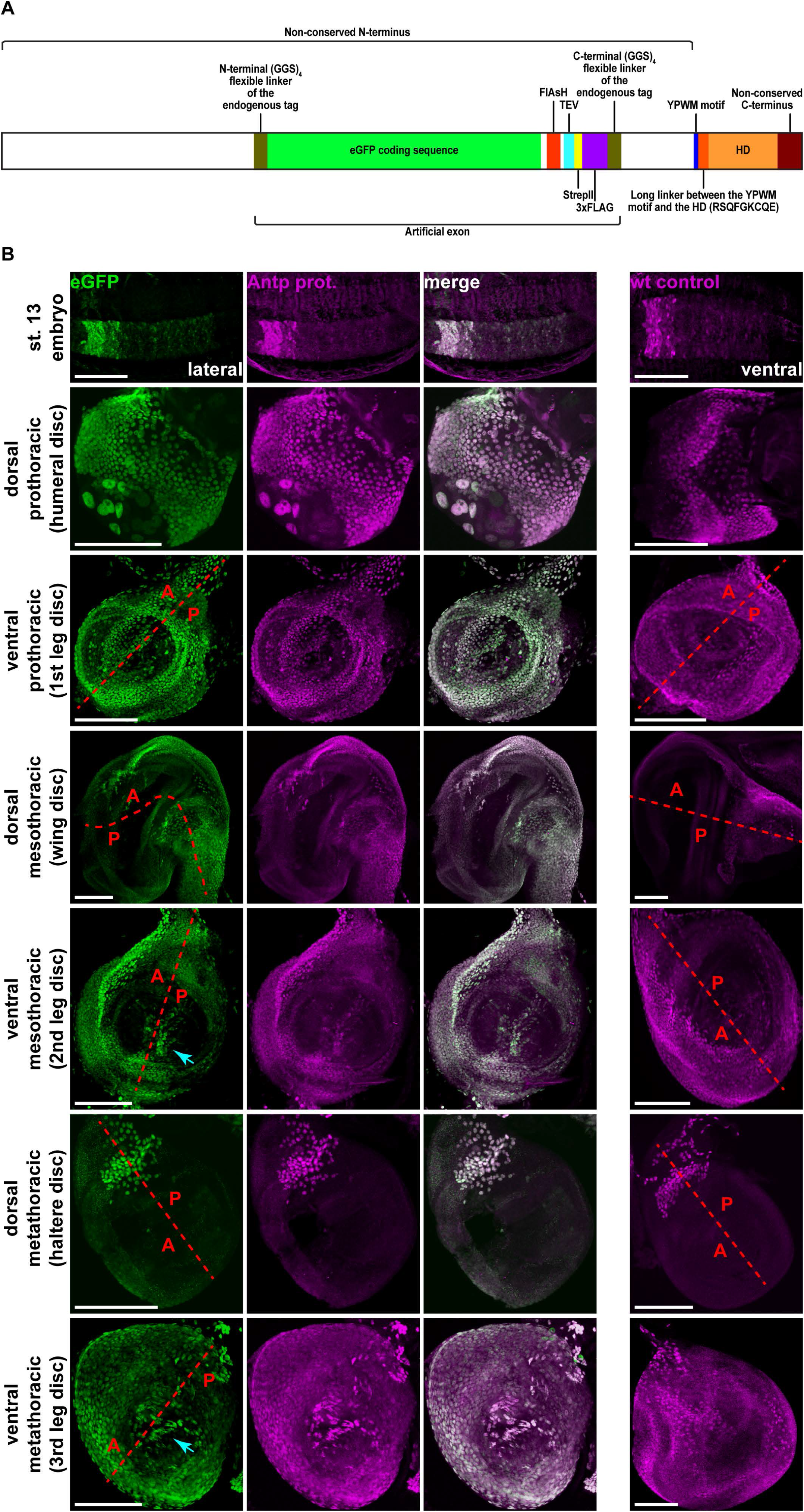
Antp expression patterns are not altered by the MiMIC MI02272 insertion. (A) Schematic representation of the Antp-eGFP fusion protein produced by the conversion of the MiMIC MI02272 construct to an artificial exon. The eGFP-encoding artificial exon is situated in intron 6 of the mRNA and is spliced in between exons 6 and 7 that correspond to the long and non-conserved N-terminal coding sequence of the protein, which has little (if any) function *in vivo* (Papadopoulos et al., 2011), and does not disrupt the homeodomain or YPWM motif. All features have been drawn to scale. (B) Heterozygous flies (embryos and third instar larvae), examined for their Antp-eGFP pattern (detected by an antibody to GFP, green), as compared to the total amount of Antp (expressed by the sum of the MiMIC *Antp-eGFP* and the wild type *Antp* loci), detected by an Antp antibody (magenta). Comparisons of the Antp pattern in wild type embryos and all thoracic imaginal discs are provided case-wise in the right panel. In discs, dashed lines approximately separate the anterior (indicated by “A”) from the posterior (indicated by “P”) domain of the disc. Note the high expression of Antp in the humeral disc. In the leg discs, Antp is expressed most strongly in the posterior compartment of the prothoracic leg disc, the anterior compartment of the mesothoracic leg disc and in an abundant pattern in the metathoracic leg disc. Cyan arrows point to Antp positive cells in the second and third leg discs that are centrally located, as previously shown (Engstrom et al., 1992). All images represent Z-projections. Scale bars denote 100 *μm.*

**Supplemental Figure S7:**
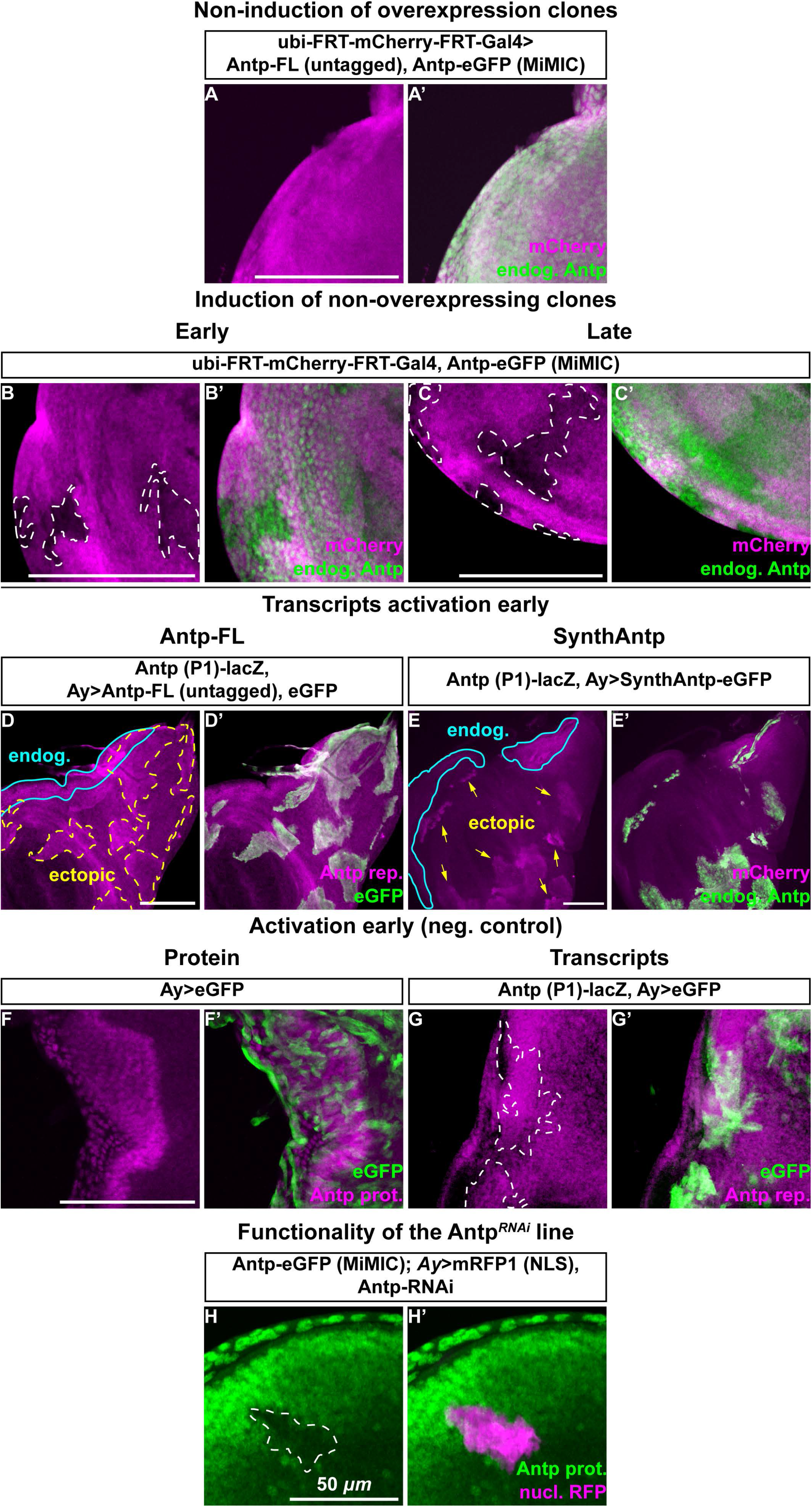
Antp is sufficient and required to trigger a developmental switch from transcriptional auto-activation to auto-repression – controls. (A-C’) Negative controls of Antp clonal auto-activation and repression using early and late clone induction regimes. (A-A’) Without induction of clones expressing full-length untagged Antp, no repression or activation of endogenous Antp protein is observed. (B-C’) Upon induction of non-overexpressing clones (clones expressing only Gal4, without a UAS transgene), no activation or repression of Antp protein is observed at early or late induction time points. White dashed lines in (B) and (C) outline the induced clones, marked by the absence of mCherry. (D-E’) Early ectopic induction of either Antp full-length untagged protein (D-D’) or SynthAntp (E-E’) result in upregulation of the Antp P1 reporter. Yellow dashed lines in (D) and arrows in (E) point to the induced clones and cyan continuous lines show the regions of high endogenous expression of the reporter. Clones have been marked by cytoplasmic eGFP. (F-G’) Negative controls of early clonal induction of eGFP alone (without concurrent induction of Antp) show no repression of the Antp protein (F-F’) or the P1 reporter (G-G’). Dashed lines in (G) mark the clones of eGFP induction. (H-H’) Positive control of clonal knockdown of the Antp *RNAi* line used in Fig. 3. Clonal knockdown by *RNAi* (indicated by the dashed line in (H) and marked by nuclear mRFP1 in (H’)) resulted in efficient downregulation of the endogenous Antp protein. Scale bars denote 100 *μm,* unless otherwise indicated.

**Supplemental Figure S8:**
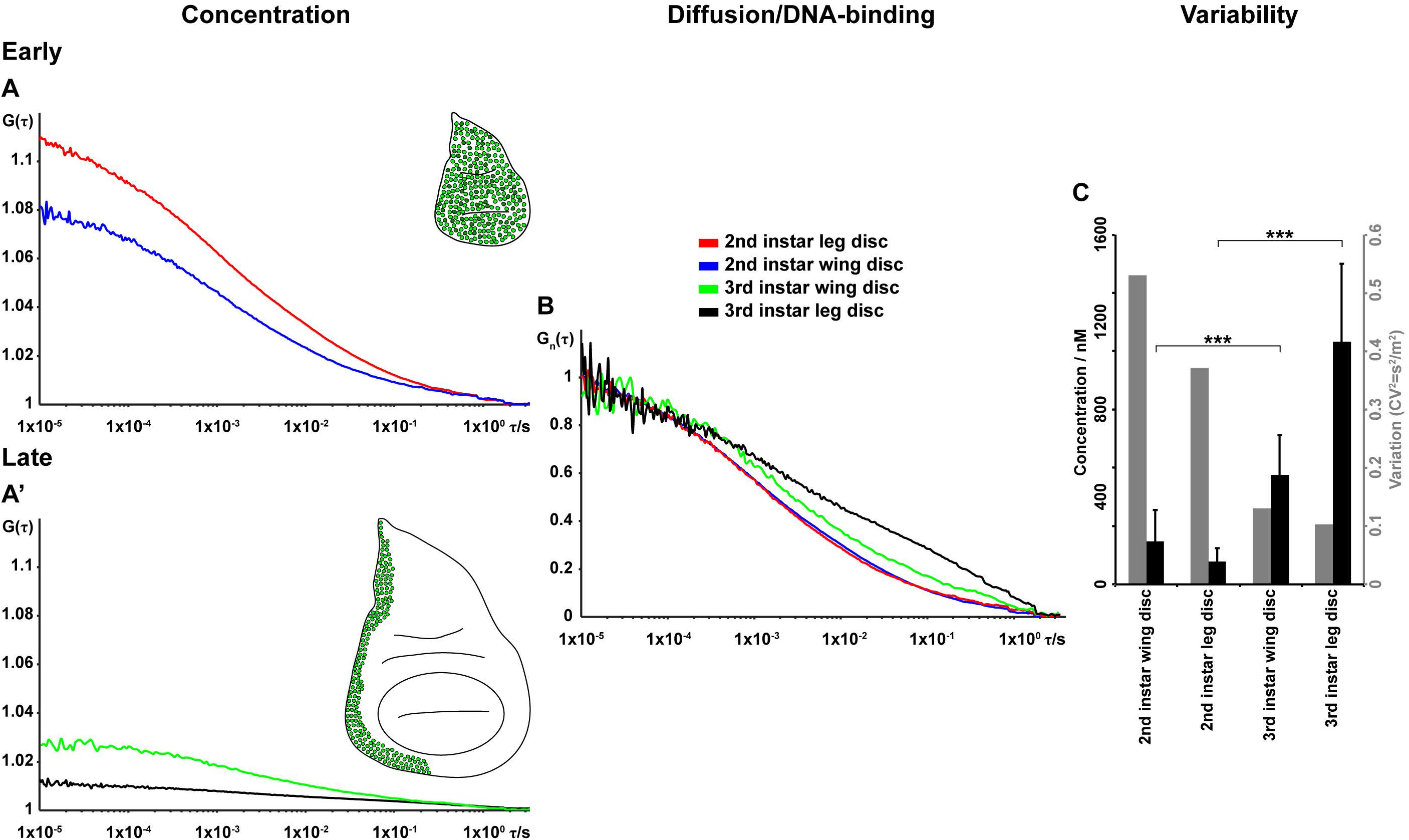
Antp concentration and cell-to-cell variability in second and third instar wing and leg imaginal discs. (A-A’’) Representative FCS curves recorded in second and third instar wing and leg imaginal discs, expressing *Antp-eGFP.* Note the low concentration in second instar leg and wing discs, reflected by the relatively high amplitude of the FCS curves (inversely proportional to concentration) in (A), as compared to the high concentration in third instar discs in (A’). (B) FCS curves shown in (A) and (A’), normalized to the same amplitude, *G_n_*(*τ*) = 1 at *τ* = 10 μ*s*, show a shift towards longer decay times in the third instar leg and wing discs, indicative of pronounced interactions of Antp with chromatin. FCS curves are color-coded as outlined in panel (A). (C) Quantification of average concentrations and cell-to-cell variability in protein concentration among neighboring nuclei in wing and leg, second and third instar, discs. Black bars denote the average concentration and grey bars denote the variability, expressed as the variance over the squared mean. Note the increase in average concentration from second to third instar (eleven-fold increase in the leg disc) and the concurrent drop in variability to almost half of its value. Statistical significance was determined using Student’s two-tailed T-test (***, p<0.001, namely *p*_3_*_rd_*_-2_*_nd instar leg_* = 4.4 · 10^−18^ and *p*_3_*_rd_*_–2_*_nd instar wing_* 3.2 · 10^−8^).

**Supplemental Figure S9:**
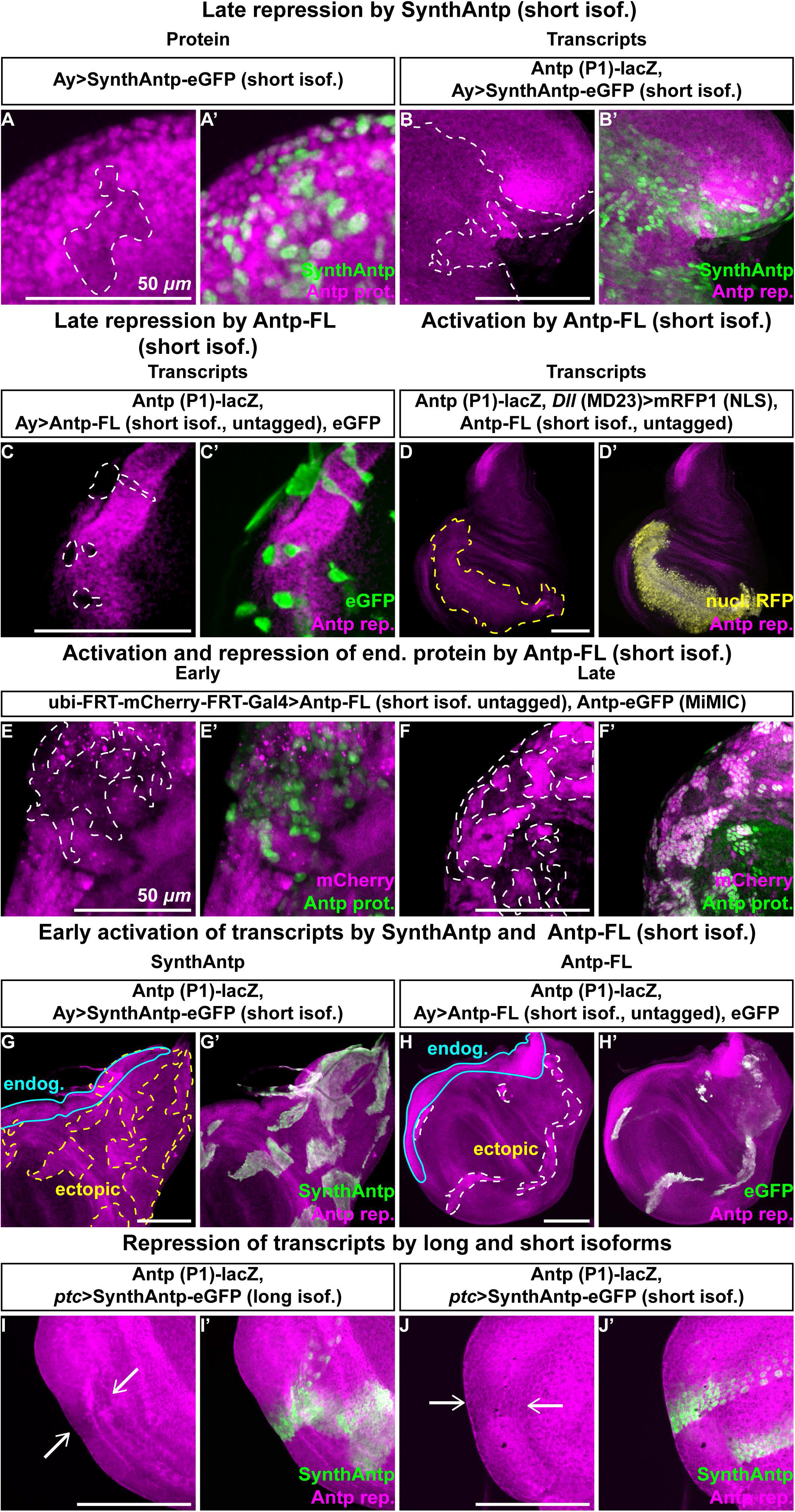
Developmental control of Antp auto-activation and repression relies on the relative concentrations of preferentially auto-activating and auto-repressing Antp isoforms, which display different binding affinities to chromatin – short linker isoform controls. (A-H) Experiments of Figs. 2–4, performed with short linker (preferentially auto-repressing) full-length and SynthAntp isoforms on their capacity to repress and activate Antp P1 reporter transcription and Antp protein. Dashed lines in all panels outline the clones induced or the region of ectopic expression using *Dll* (MD23)-Gal4, whereas closed continuous cyan lines outline the regions of endogenous Antp P1 reporter expression in (G) and (H). (A-A’) Repression of Antp protein by late clonal induction of SynthAntp in the wing notum. (B-B’) Equivalent assay as in (A-A’), but monitoring auto-repression of the Antp P1 promoter transcription. (C-C’) Similar assay to (B-B’), using the full-length Antp protein, induced at the later time point. (D-D’) Ectopic induction of full-length, short linker, untagged Antp cDNA with concurrent labeling of the expression domain by nuclear mRFP1 results in weak ectopic auto-activation of the Antp P1 reporter. (E-F’) Early and late clonal induction of full-length, short linker, untagged Antp results in auto-repression (E-E’), or induction (F-F’), of the endogenous Antp protein, respectively. (G-H’) Early clonal induction of SynthAntp (G-G’) or the full-length cDNA (H-H’), both featuring a short linker, triggers ectopic activation of P1 promoter transcription. (I-J’) Antp long (I-I’) and short (J-J’) linker isoforms repress Antp at the transcriptional level (monitored by Antp P1 reporter expression) when induced by *ptc-*Gal4 in the wing disc. Arrows point to the regions of auto-repressed Antp promoter. Scale bars denote *100 μm,* unless otherwise indicated.

**Supplemental Figure S10:**
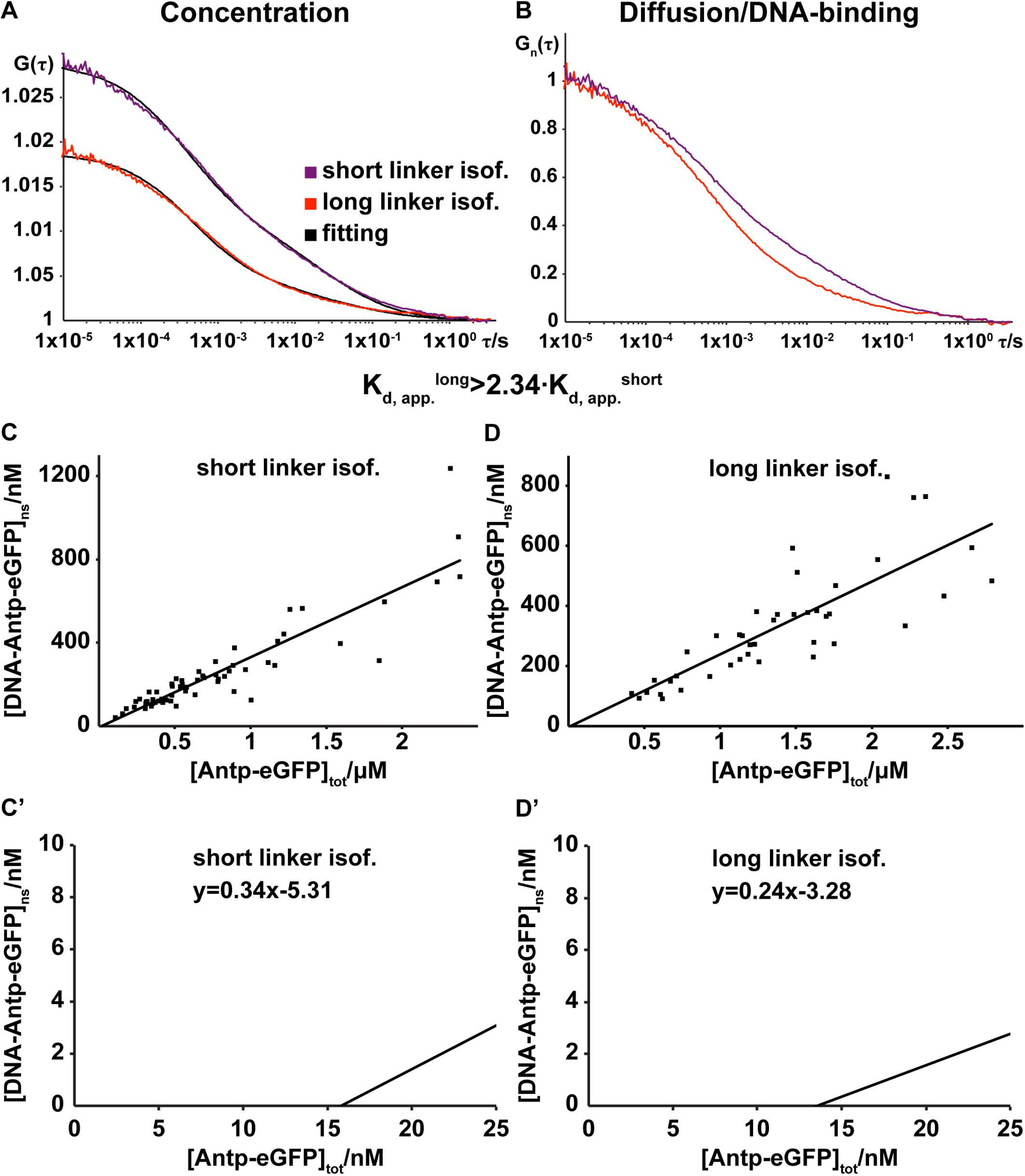
Comparative binding study of Antp short and long linker isoforms by FCS. (A-B) FCS analysis performed on third instar wing and antennal imaginal discs, expressing short or long linker Antp isoforms (tagged to eGFP) close to endogenous concentrations, from the *69B*-enhancer. Cell nuclei of similar concentrations in the two datasets have been selected for analysis (A). Average FCS measurements on the short linker Antp isoform display a consistent shift towards longer decay times, as compared to its long linker counterpart (B), indicating higher degree of chromatin binding. (C-D’) Binding study of short and long linker Antp isoforms in third instar wing and antennal discs, expressed by *69B*-Gal4. The concentration of the Antp short and long linker isoform DNA-bound complexes (derived by fitting the FCS curves in (A)) is plotted as a function of the total concentration of Antp-eGFP molecules. From the linear regression equations, *y =* 034*x−*531 (D’) and *y =* 0.24*x−*3.28 (E’), the ratio of apparent dissociation constants for the long and short linker isoforms was calculated to be 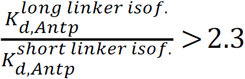 (for the calculation refer to Supplement 3). The two dissociation constants differ at least 2.3 times, indicating stronger binding of the short linker isoform to the DNA, as compared to the long linker one.

**Supplemental Figure S11:**
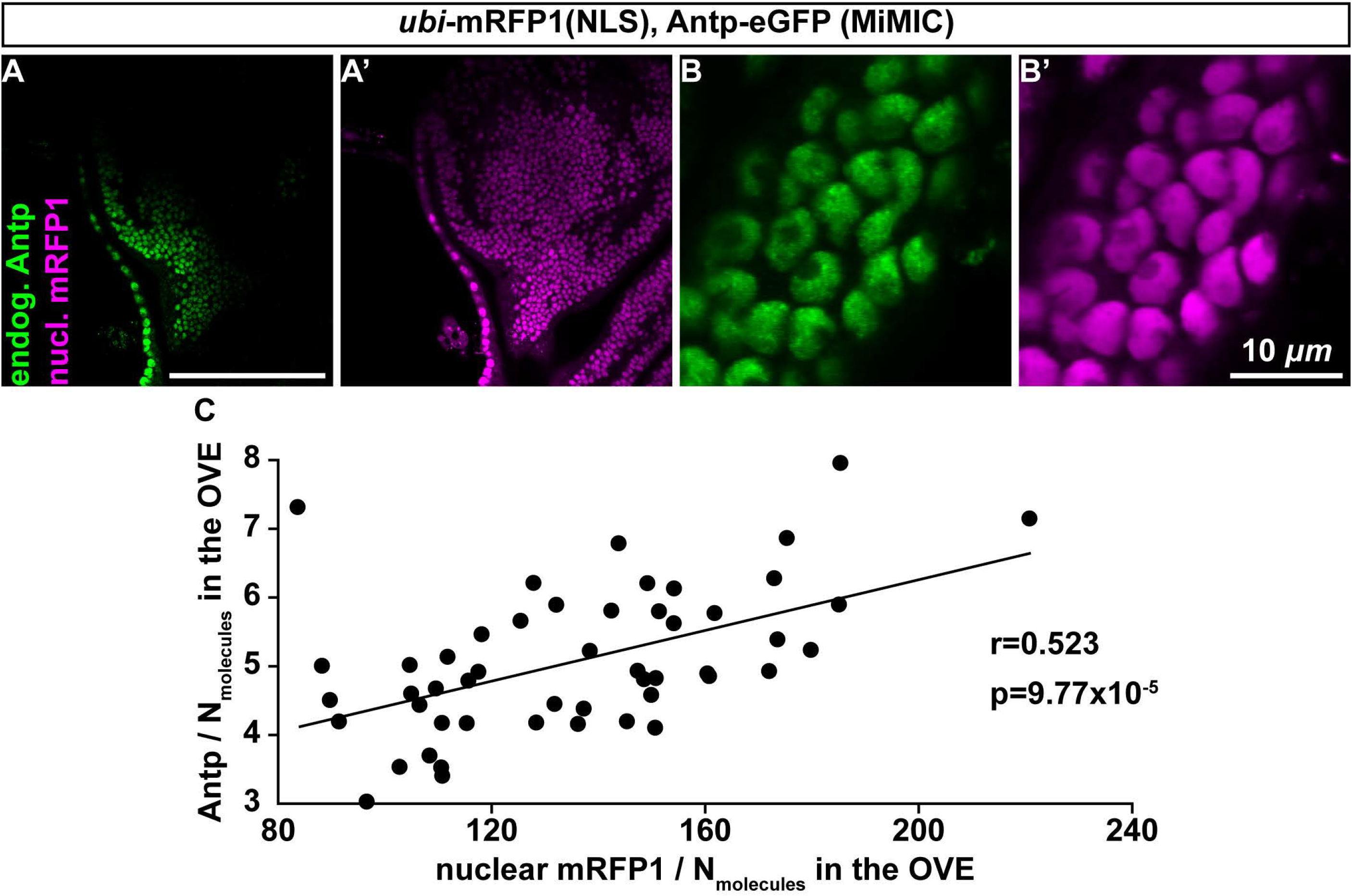
Investigation of extrinsic variability in the endogenous Antp-eGFP expression domain. (A-B’) Live imaging of a wing disc notum, where nuclear mRFP1 protein is highly expressed from a constitute enhancer (*ubi*-mRFPI(NLS)), alongside with endogenous Antp-eGFP. FCS measurements were performed in the region of high co-expression of Antp and mRFP1. (B-B’) Higher magnification of cells as in (A-A’). Note the uneven distribution of Antp in the nuclei and the formation of sites of accumulation in (B). (C) Plot of the concentration of Antp-eGFP (expressed as number of fluorescent molecules in the Observation Volume Element (OVE)). The correlation coefficient *r* was calculated to be *r = 0.523* and the p-value to be *p_correlation_ =* 9.77 · 10^−5^. Scale bars denote 100 *μm,* unless otherwise indicated.

**Supplemental Figure S12:**
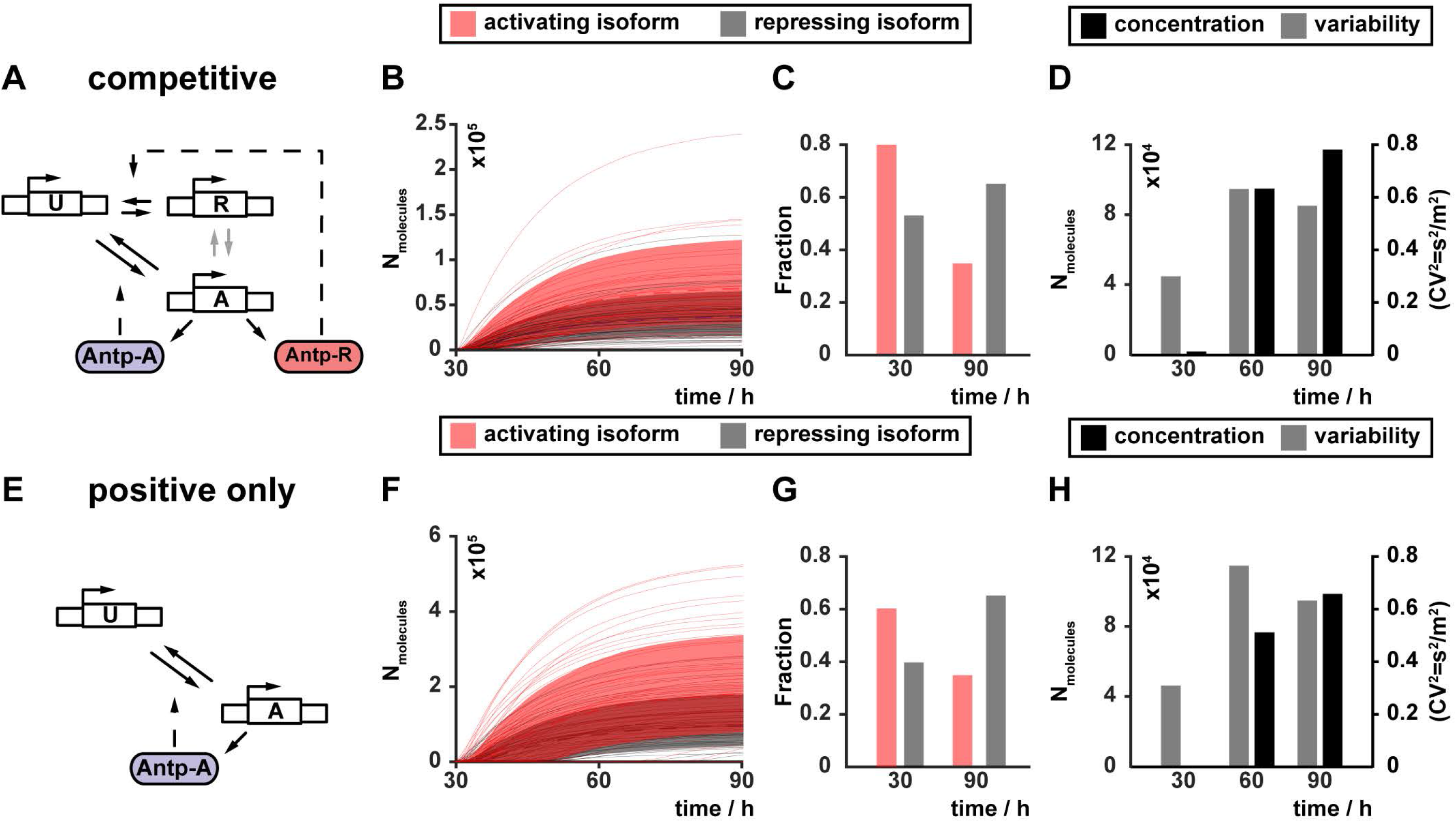
(A-D) Competition of Antp binding, whereby state “A” can be reached only through the unbound state “U” in (A), results in increase in Antp protein numbers (D) without decrease in variability (grey bars in (D)). Trajectories of individual simulations are presented in (B) and the distribution of the Antp isoforms, predicted by the model, in (C). (E-H) Requirement of the negative feedback for suppression of variability. In the absence of the state “R” (E), concentration increases (H), but variability also increases rather than being suppressed (grey bars in (H)). Trajectories of individual simulations are presented in (F) and the distribution of the Antp isoforms, predicted by the model, in (G).

**Supplemental Figure S13:**
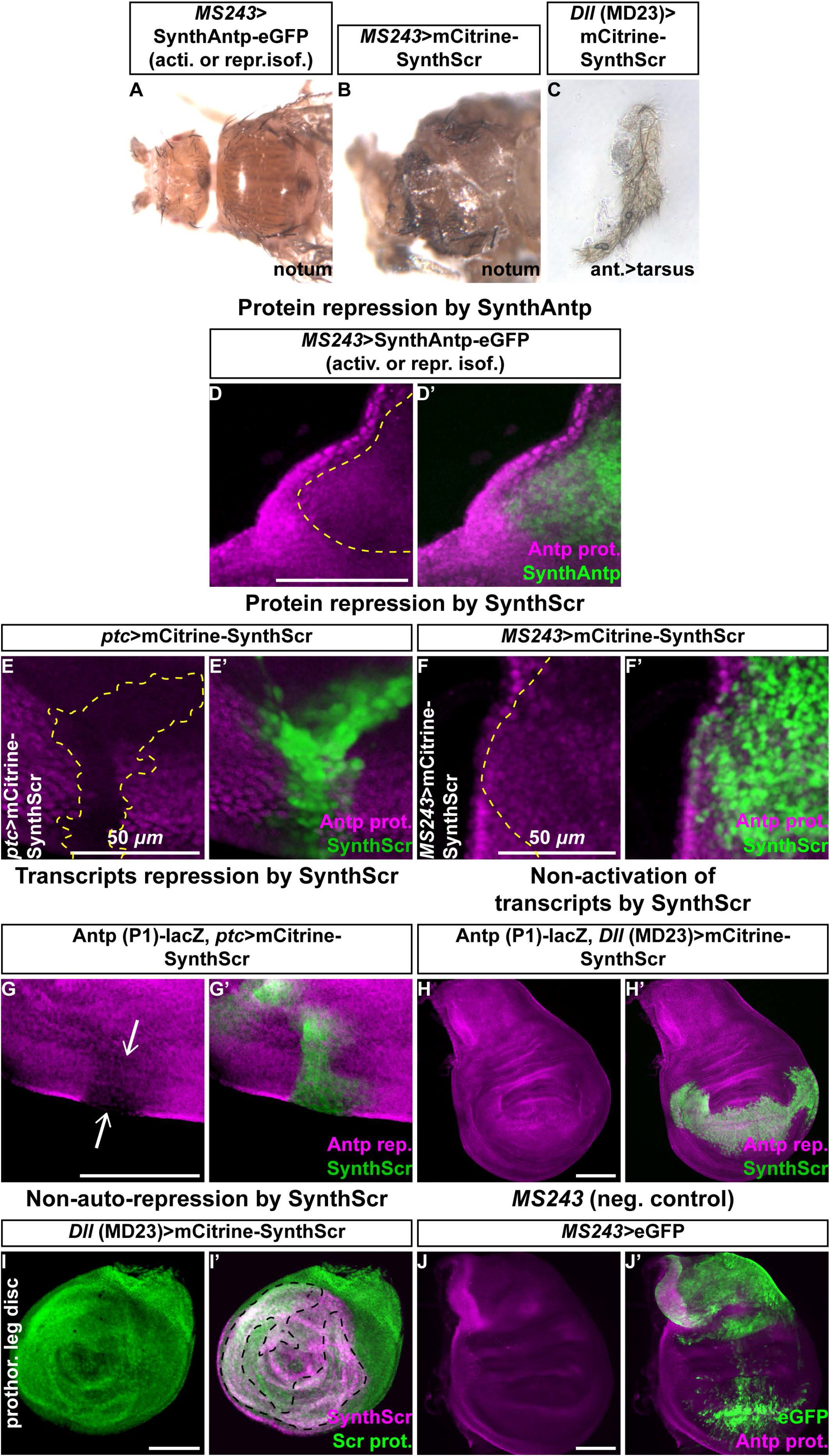
Controls of Antp model predictions and Scr-mediated perturbations. (A-C) Perturbations of the model system in (Fig. 5A) by overexpression of Antp long or short linker isoforms or an exogenous Antp repressor (Scr). Overexpression of activating or repressing SynthAntp-eGFP isoforms by *MS243*-Gal4 results in normal development of the fly notum (A), whereas induction of an exogenous repressor (mCitrine-SynthScr) results in severe malformations, indicated by developmental defects of the adult cuticle (B). Flies of both genotypes in (A) and (B) die as pharate adults. (C) Induction of mCitrine-SynthScr in the antennal disc results in complete transformations of antenna to tarsus as the induction of SynthAntp-eGFP (Fig. 6A). (D-D’) *MS243*-Gal4-mediated expression of repressing or activating SynthAntp isoforms results in repression of the Antp endogenous protein in the notum region of the wing disc. (E-E’) Ectopic expression of SynthScr in the wing disc using *ptc-*Gal4 results in drastic reduction of endogenous Antp protein levels. (FF’) Ectopic expression of SynthScr by *MS243*-Gal4 in the notum results in repression of the Antp protein. (G-G’) SynthScr represses Antp at the transcriptional level, as indicated by the absence of transcription of the Antp P1 reporter (white arrows in (G)). (H-H’) Unlike SynthAntp, SynthScr is not able to activate the Antp P1 promoter reporter transcription (H), when induced by *Dll* (MD23)-Gal4. (I-I’) SynthScr is not able to downregulate its own endogenous protein levels upon overexpression by *Dll* (MD23)-Gal4. Dashed line in (I’) outlines the region of high overlap between the overexpressed SynthScr and endogenous Scr stainings. (J-J’) Negative control staining for the induction of eGFP in the wing disc notum by *MS243*-Gal4, which fails to repress endogenous Antp protein. Dashed lines in (D), (E) and (F) outline the regions of ectopic overexpression of SynthAntp or SynthScr, where endogenous Antp is repressed, whereas the dashed line in (I’) outlines the region of overlap between SynthScr overexpression and endogenous expression of the Scr protein, where no repression is observed. Scale bars denote 100 *μm,* unless otherwise indicated.

## Supplemental information

### Supplement 1: Background on Fluorescence Microscopy Imaging and FCS

Two individually modified instruments (Zeiss, LSM 510 and 780, ConfoCor 3) with fully integrated FCS/CLSM optical pathways were used for imaging. The detection efficiency of CLSM imaging was significantly improved by the introduction of APD detectors. As compared to PMTs, which are normally used as detectors in conventional CLSM, the APDs are characterized by higher quantum yield and collection efficiency – about 70 % in APDs as compared to 15 – 25 % in PMTs, higher gain, negligible dark current and better efficiency in the red part of the spectrum. Enhanced fluorescence detection efficiency enabled image collection using fast scanning (1–5*/pixel*). This enhances further the signal-to-noise-ratio by avoiding fluorescence loss due to triplet state formation, enabling fluorescence imaging with single-molecule sensitivity. In addition, low laser intensities (150–750 *μW*) could be applied for imaging, significantly reducing the photo-toxicity (Vukojevic et al., 2008).

FCS measurements are performed by recording fluorescence intensity fluctuations in a very small, approximately ellipsoidal observation volume element (OVE) (about 0.2*μm* wide and 1μ*m* long) that is generated in imaginal disc cells by focusing the laser light through the microscope objective and by collecting the fluorescence light through the same objective using a pinhole in front of the detector to block out-of-focus light. The fluorescence intensity fluctuations, caused by fluorescently labeled molecules passing through the OVE are analyzed using temporal autocorrelation analysis.

In temporal autocorrelation analysis we first derive the autocorrelation function

*G*(τ):

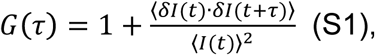

where *δI*(*t*) = *I*(*t*) – 〈*I*(*t*)〉 is the deviation from the mean intensity at time *t* and *δI*(*t* + *τ*) = *I*(*t + τ*) -〈*I*(*t*)〉 is the deviation from the mean intensity at time *t* + τ. For further analysis, an autocorrelation curve is derived by plotting *G*(*τ*) as a function of the lag time, i.e. the autocorrelation time *τ*.

To derive information about molecular numbers and their corresponding diffusion time, the experimentally obtained autocorrelation curves are compared to autocorrelation functions derived for different model systems, and the model describing free three dimensional (3D) diffusion of two components and triplet formation was identified as the simplest and best suited for fitting the experimentally derived autocorrelation curves, and was used throughout:

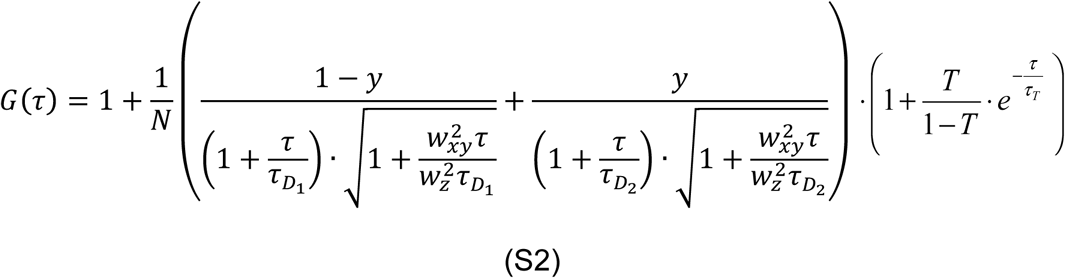

In the above equation, *N* is the average number of molecules in the OVE; *y* is the fraction of the slowly moving Antp-eGFP molecules; 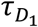 is the diffusion time of the free Antp-eGFP molecules; 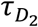 is the diffusion time of Antp-eGFP molecules undergoing nonspecific interactions with the DNA; *w_xy_* and *w_z_* are radial and axial parameters, respectively, related to spatial properties of the OVE; T is the average equilibrium fraction of molecules in the triplet state; and τ*_Τ_* the triplet correlation time related to rate constants for intersystem crossing and the triplet decay. Spatial properties of the detection volume, represented by the square of the ratio of the axial and radial parameters 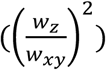, are determined in calibration measurements performed using a solution of Rhodamine 6G for which the diffusion coefficient (D) is known to be *D_Rh_*_6_*_G_* = 4.1 · 10^−10^ *m*^2^*s*^−1^ (Muller et al., 2008). The diffusion time, *τ_D_*, measured by FCS, is related to the translation diffusion coefficient *D* by:

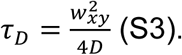

To establish that Antp molecules diffusing through the OVE are the underlying cause of the recorded fluorescence intensity fluctuations, we plotted the characteristic decay times 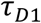 and 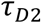, obtained by FCS, as a function of the total concentration of Antp molecules (Supplemental Fig. S2). We observed that both characteristic decay times remain stable for increasing total concentration of Antp molecules, signifying that the underlying process triggering the fluorescence intensity fluctuations is diffusion of fluorescent Antp molecules through the OVE (which is independent of the total concentration of Antp molecules).

In order to ascertain that the interpretation and fitting of FCS curves is correct, we have: (1) tested several laser intensities in our FCS measurements and have utilized the highest laser intensity, for which the highest counts per second and molecule (CPSM) were obtained, while photobleaching was not observed; (2) we have established that CPSM do not change among FCS measurements performed in cells expressing Antp endogenously, or overexpressed with different Gal4 drivers. Moreover, we have previously shown that both characteristic decay times increase when the size of the OVE is increased (Fig. 4 in (Vukojevic et al., 2010)). Together, these lines of evidence indicate that both short and long characteristic decay times are generated by molecular diffusion rather than by photophysical and/or chemical processes such as eGFP protonation/deprotonation; (3) we have ascertained that the long characteristic decay time of our FCS measurements is not the result of photobleaching and that differences in the relative amplitudes of the fast and slow diffusing components reflect differences in their concentrations among cells.

While we have taken all possible precautions to ascertain that the correct model for FCS data fitting is applied, some inevitable limitations still remain. For example, FCS cannot account for Antp molecules with irreversibly photobleached fluorophores or with fluorophores residing for various reasons in dark states. In addition, FCS cannot account for Antp molecules associated with large immobile structures, such as specifically bound Antp molecules. These molecules contribute to the overall background signal, but they do not give rise to fluorescence intensity fluctuations. As a consequence, transcription factor concentration can be somewhat underestimated by FCS.” In contrast, the number of transcription factor molecules may also be overestimated by FCS, when high background signal as compared to fluorescence intensity may lead to an artificially low amplitude of FCS curves, and, hence, overestimation of molecular numbers. To avoid artifacts due to photobleaching, the incident laser intensity was kept as low as possible but sufficiently high to allow high signal-to-noise ratio. This is because photobleaching of fluorophores may induce errors in the measurements of molecular numbers and lateral diffusion, yielding both smaller number of molecules and shorter values of τ_D_, and hence apparently larger diffusion coefficients. Finally, contribution of brightness, *i.e.* brightness squared, to the correlation function was not analyzed, which may in turn affect quantification of Antp numbers.

### Supplement 2: Calculation of the concentration of endogenous TFs and average number of molecules in imaginal disc cell nuclei from FCS measurements (exemplified for Antp)

Experimentally derived FCS curves were analyzed by fitting, using the model function for free three-dimensional diffusion of two components with triplet formation, equation (S2), to derive the average number of molecules in the OVE (*N*); the diffusion time of the free Antp-eGFP molecules 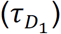; the diffusion time of Antp-eGFP molecules undergoing interactions with the DNA 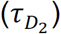; and the relative fraction of Antp-eGFP molecules that are engaged in interactions with chromatin and therefore move slowly (*y*).

In order to translate the average number of molecules in the OVE (*N*) into molar concentration, the size of the OVE, *i.e.* the axial and radial parameters (*w_z_* and *w_xy_*, respectively) were determined in calibration experiments with Alexa488 or Rhodamine 6G dyes, using equation (S3). The volume of the OVE, approximated by a prolate ellipsoid, was determined as follows:

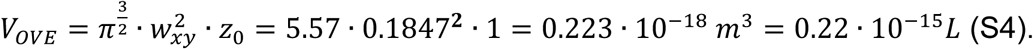

Thereafter, the average number of molecules in the OVE (*N*) was converted into molar concentration (*C*) using the relationship:

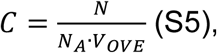

where *N_A_* is the Avogadro number (6.022 · 10^23^*mol*^−1^), which indicated that one molecule in the OVE corresponds on the average to 8.74 *nM* concentration of fluorescent molecules in the nucleus.

Finally, the concentration of non-specifically bound TF molecules ([*DNA − Antp – eGFP*]_ns_) was calculated by multiplying the relative amplitude of the second component (*y*), determined by fitting the experimental autocorrelation curves with the model function (S2), with the total concentration of Antp-eGFP, *C =* [*Antp – eGFP*]_0_, which was determined from the amplitude of the autocorrelation curve at zero lag time:

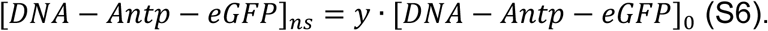

The concentration of non-specifically bound TF molecules ([*DNA – Antp – eGFP*]*_ns_*) was then plotted as a function of the total concentration of Antp-eGFP ([*Antp – eGFP*]_0_) to yield the graphs shown in_Supplemental Fig. S10C,D.

In order to estimate the total number of molecules in the wing disc imaginal cell nuclei, we applied the following calculation. The wing disc nuclei within the Antp expression domain (prescutum precursors) are not spherical, but rather ellipsoidal. Their axes were determined by fluorescence imaging to be 1.4*μm* in the transverse dimension and 2.8 *μm* in the longitudinal. The volume of the nucleus was approximated by the volume of a prolate ellipsoid:

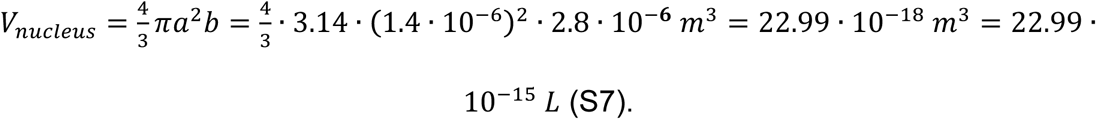

Therefore, the OVE represents roughly 1/121 of the nuclear volume:

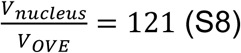

and the number of molecules in Antp-eGFP nuclei is on the average 57.37 · 121 *≈* 6942 molecules in third instar wing and 127 · 121 ≈ 15367 in third instar leg discs.

Generalizing, with known axial and radial parameters of the OVE and calculation of the transverse and longitudinal dimensions of the nucleus, the total number of molecules of transcription factor in the nucleus can be estimated:

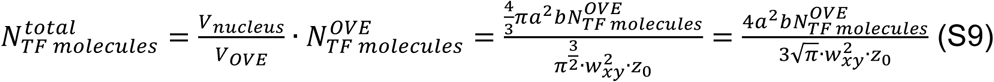

### Supplement 3: Calculation of the ratio of apparent Antp dissociation constant for short and long linker Antp isoforms from FCS measurements on ectopically expressed Antp

Antp undergoes both specific and non-specific interactions with DNA, with nonspecific interactions preceding the specific ones and effectively assisting the binding to a specific target site by facilitated diffusion (Halford and Marko, 2004). The searching for specific binding sites can be described as a two-step process of consecutive reactions (Vukojevic et al., 2010):

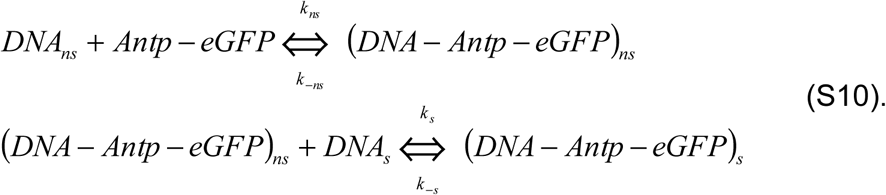

The turnover rate for the non-specific complex is:

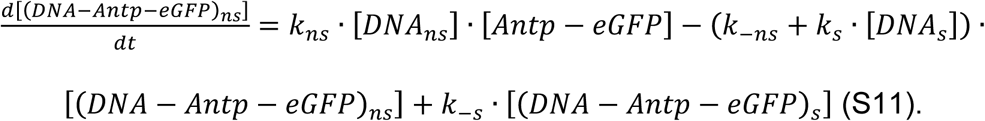

Assuming a quasi-steady state approximation:

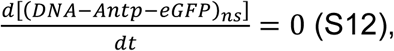

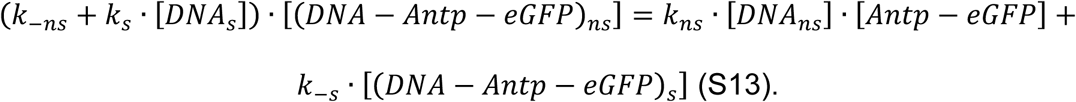

Using the mass balance equation to express the concentration of the free TF:

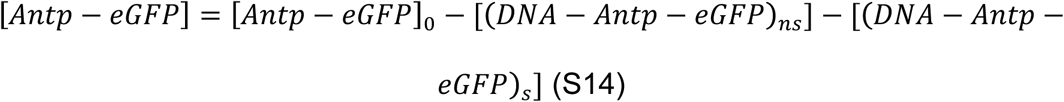

and assuming that:

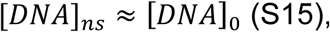

equation (S13) becomes:

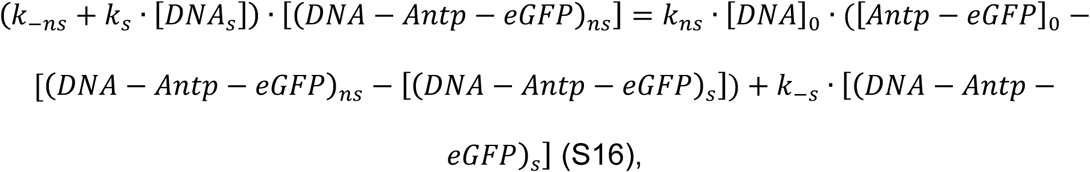

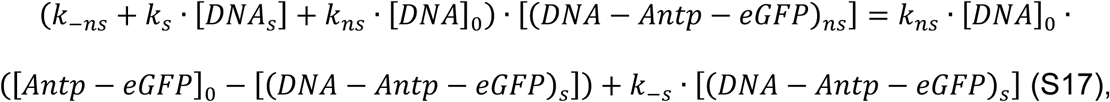

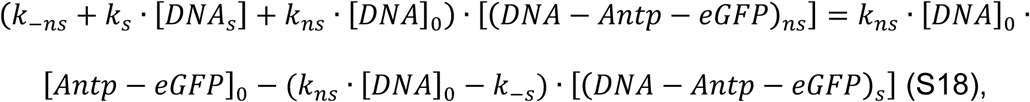

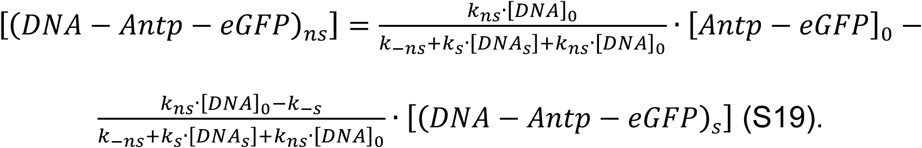

According to equation (S19) and the FCS data presented in Supplemental Fig. S10, the slope of the linear dependence for:

a) the short linker Antp isoform gives:

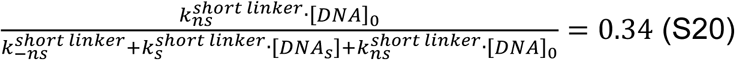

and the intercept:

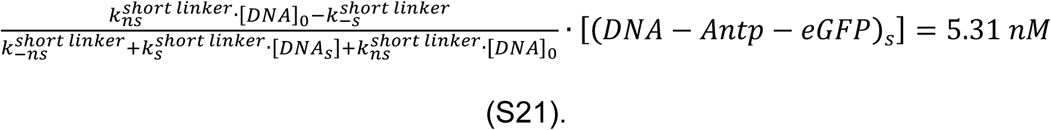

If 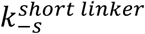 is small compared to 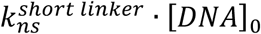 and can therefore be neglected, then:

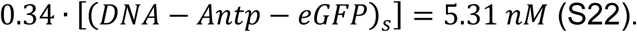

Thus, the concentration of specific complex between Antp-eGFP and DNA in the wing disc cell nuclei can be estimated to be:

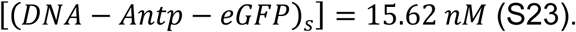

The average concentration of free-diffusing Antp-eGFP molecules is determined as follows:

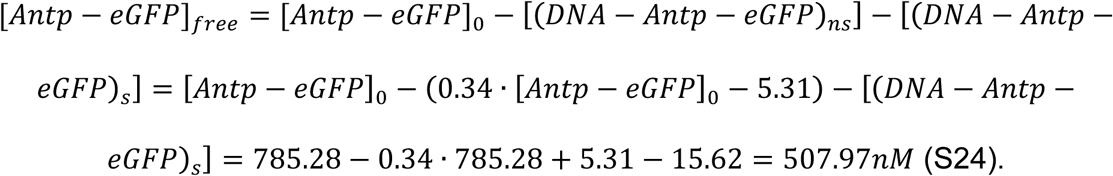

Using the experimentally determined concentration of specific DNA–Antp-eGFP complexes (equation (S23)), we could estimate the dissociation constant for the specific DNA–Antp-eGFP, as a function of the total concentration of specific Antp binding sites, to be:

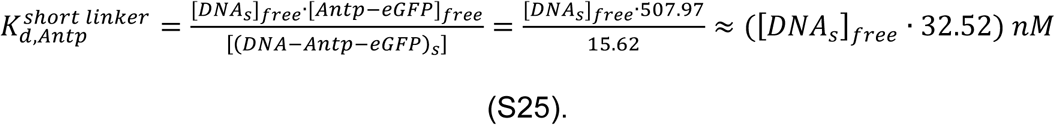

b) The long linker Antp isoform gives:

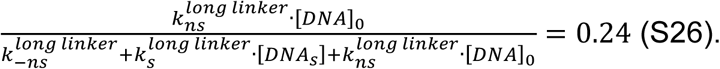

and the intercept:

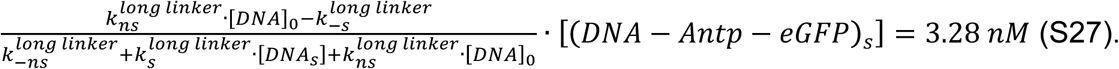

If 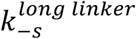 is small compared to 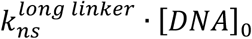 and can therefore be neglected, then:

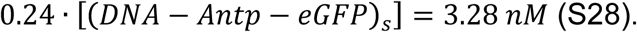

Thus, the concentration of specific complex between Antp-eGFP and DNA in the wing disc cell nuclei can be estimated to be:

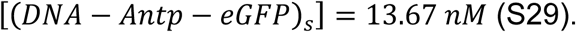

The average concentration of free-diffusing Antp-eGFP molecules is determined as follows:

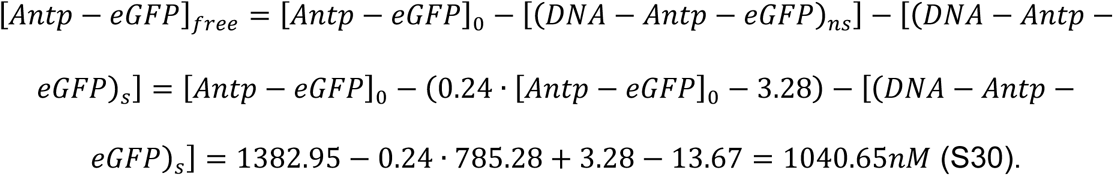

Using the experimentally determined concentration of specific DNA–Antp-eGFP complexes (equation (S29)), we could estimate the dissociation constant for the specific DNA–Antp-eGFP, as a function of the total concentration of specific Antp binding sites, to be:

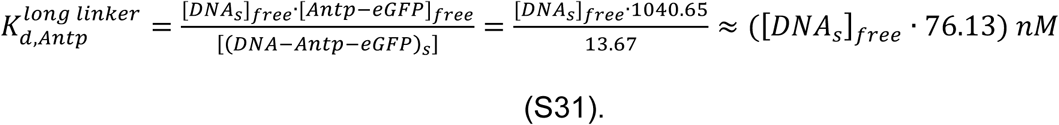

From equations (S25) and (S31), we could calculate the ratio of the apparent equilibrium dissociation constants for specific interactions to be:

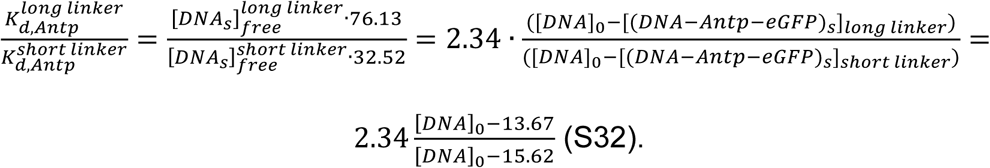

Therefore:

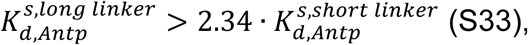

independently of the total concentration of Antp binding sites in the nucleus. Although the affinities of the ratio of the equilibrium dissociation constants does not depend on the total concentration of binding sites for Antp ([*DNA*]_0_), its value is required for the calculation. For values close to 15.62 *nM* ([*DNA*]_0_ → 15.62 *nM*) with [*DNA*]_0_ > 15.62 *nM*, the ratio of the apparent equilibrium dissociation constants will be high:

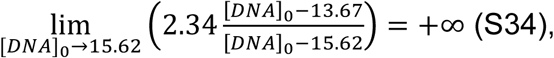

indicating that, in this case, the short linker isoform will bind the Antp binding sites with much higher affinity than the long linker isoform.

In contrast, for considerably higher values of [*DNA*]_0_ than 15.62 *nM* ([*DNA*]_0_ → +∞), the ratio of apparent equilibrium dissociation constants will be:

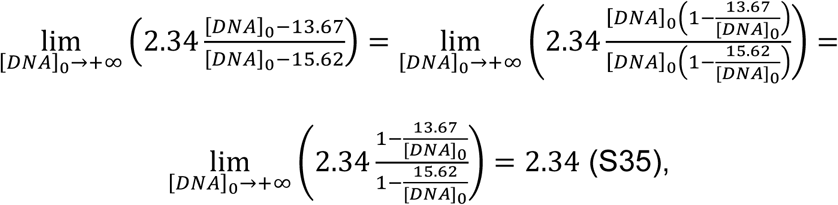

indicating a roughly 2.5fold higher affinity of the short linker repressive isoform.

In addition, equations (S20) and (S26) contain information about the ratio of the apparent equilibrium dissociation constants for nonspecific interactions [(Vukojevic et al., 2010), Fig. 5, solid red line *versus* dashed red line]. Thus, the slopes of the linear regression lines (Supplemental Fig. S10C-D’), give:

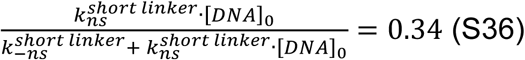

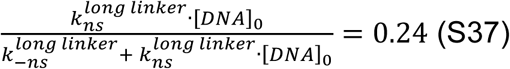

From these relationships, the ratio of the apparent equilibrium dissociation constants for nonspecific interactions can be estimated to be:

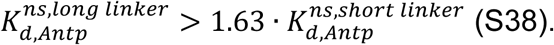

Thus, our analysis shows that the short linker, which is the preferentially repressing isoform, binds with higher affinity (lower *K_d_*) to both specific and nonspecific binding sites on the DNA [(S33) and (S38), respectively]. This, in turn, implies that the short linker is also more efficient in searching for specific TF binding sites, as evident from the lower dissociation constant for nonspecific DNA interactions of the short linker isoform (Sela and Lukatsky, 2011; Soltani et al., 2015), and that it binds with lower apparent dissociation constant to specific binding sites on the DNA.

### Supplement 4: Stochastic modeling of Antennapedia expression

In the following, we develop a simple mathematical model that is able to explain the behavior of Antennapedia (Antp) expression at early and late developmental stages. The Antp promoter is modeled as a continuous-time Markov chain with three distinct transcriptional states. In the absence of Antp, the promoter is in an unbound state (“U”), in which transcription is inactive. From this state, the promoter can switch to a transcriptionally active state “A” at a rate, assumed to be proportional to the concentration of the long-linker, activating isoform of Antp. Analogously, repression of the promoter by the short-linker isoform of Antp is modeled by an additional transcriptionally inactive state “R”, which can be reached from state “U” at a rate proportional to the concentration of that isoform. The corresponding reverse transitions from states “R” and “A” back into state “U” are assumed to happen at a constant rate *k.* Since the activating isoform can potentially also repress the promoter, we assume that state “R” can be reached also from the active state “A”. Similarly, we model a potential link also in the reverse direction from state “A” to “R”. Depending on the model variant, we assume this transition happens either at a constant rate *k* (competitive promoter model) or at a rate proportional to the concentration of the repressing isoform of Antp (non-competitive promoter model). In the latter case, repression through short-linker isoforms can take place even if a long-linker isoform is already bound to the promoter. As we have demonstrated in Fig. 5A,B, the two model variants yield qualitative differences in Antp expression. For the sake of illustration, the following description focuses on the non-competitive model variant but we remark that the competitive model can be derived analogously.

At a particular time point *t,* the transcription rate of Antp is determined by the current state of the promoter, i.e., *λ*(*t*)*∊*[0*, λ_A_,* 0], with *λ_A_* as the transcription rate associated with state “A”. In line with our experimental findings, we assume that transcripts are spliced into the activating and repressing isoforms at different rates *ρ_A_* and *ρ_R_*, respectively. This allows us to capture the imbalance between the two isoforms that was revealed by our FCS data. The overall expression rates for the two isoforms of Antp are then given by *h_A_*(*t*) *= λ*(*t*)*Zρ_A_* and *h_R_*(*t*) *= λ*(*t*)*Zρ_R_*, whereas *Z* is a random variable that accounts for extrinsic variability in gene expression rates (Zechner et al., 2012). In all of our analyses, we model *Z* as a Gamma random variable, i.e., *Z~Γ*(*α, β*) with *α* and *β* as shape and inverse scale parameters of that distribution. In summary, we describe the auto-regulatory circuit of Antp expression by a Markovian reaction network of the form:

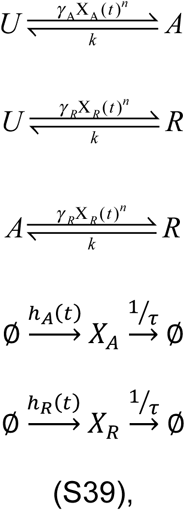

with *X_A_*(*t*)and *X_R_*(*t*) as the concentration of the activating and repressing isoforms of Antp, *τ* as the protein half-live and *n* as a coefficient accounting for cooperativity in the binding of Antp to the promoter. The initial conditions *X_A_*(0) and *X_R_* (0) were drawn randomly in accordance with our concentration measurements at early stages. In particular, we assume that the total amount of Antp *X_tot_* in each cell is drawn from a negative binomial distribution such that 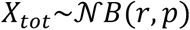, with *μ_X_* = *r*(1 − *p*)/*p* and 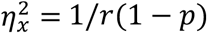 as the mean and squared coefficient of variation of this distribution. The total number of Antp molecules was then randomly partitioned into fractions of repressing and activating isoforms according to a Beta distribution. More specifically, we set *X_q_*(0) = *WX_tot_* and *X_R_*(0) = (1 – *W*)*X_tot_* with *W*~*Beta*(*a*, *b*). The parameters *r*, *p*, *a* and *b* were chosen based on our experimental data (see Table 1).

**Table 1:**
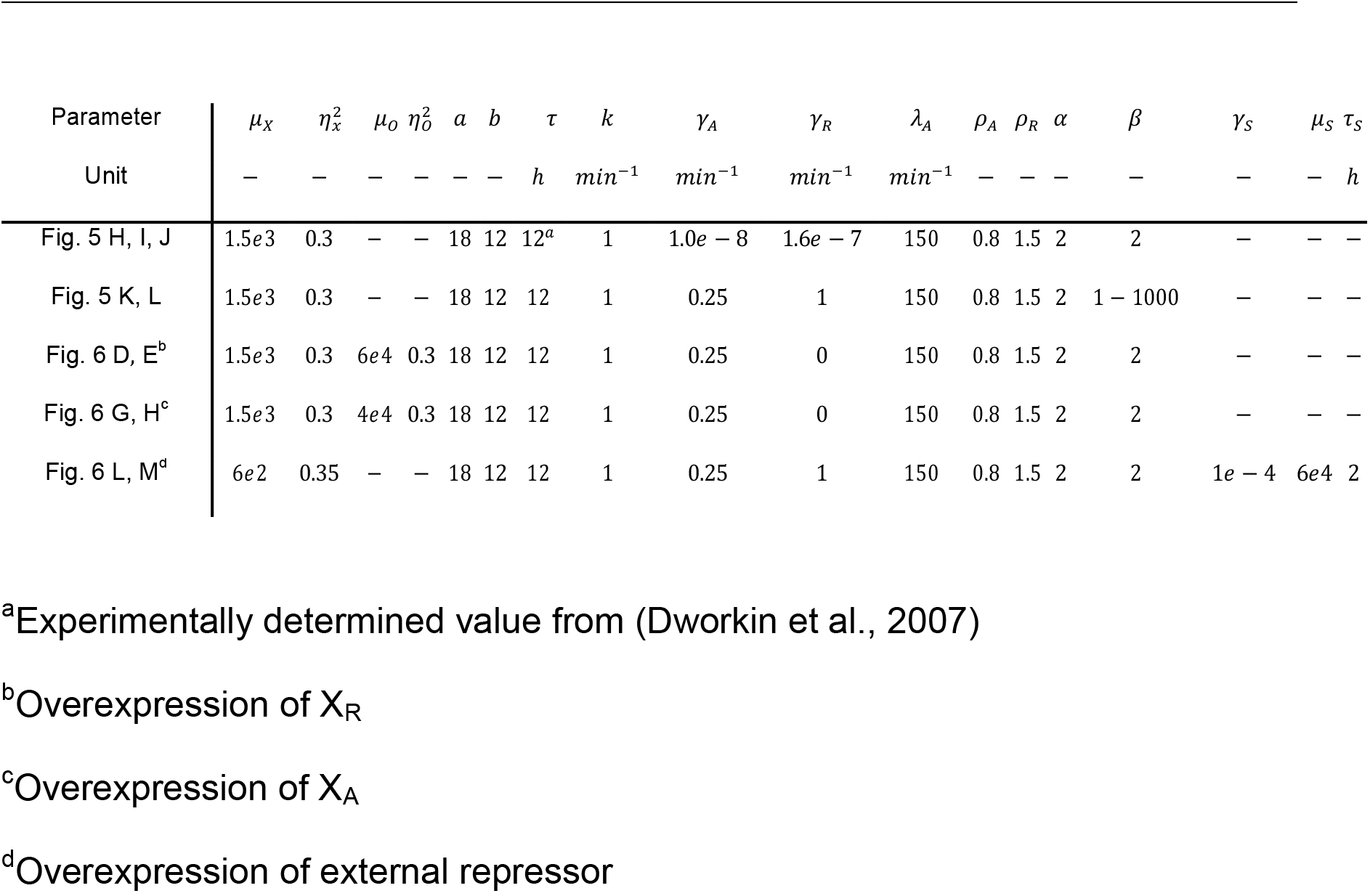
Parameters used for simulating the stochastic model of Antp expression

| Parameter | $\mu_X$ | $\eta_X^2$ | $\mu_O$ | $\eta_O^2$ | $a$ | $b$ | $\tau$ | $k$ | $\gamma_A$ | $\gamma_R$ | $\lambda_A$ | $\rho_A$ | $\rho_R$ | $\alpha$ | $\beta$ | $\gamma_S$ | $\mu_S$ | $\tau_S$ |
| --- | --- | --- | --- | --- | --- | --- | --- | --- | --- | --- | --- | --- | --- | --- | --- | --- | --- | --- |
| Unit | – | – | – | – | – | – | $h$ | $\text{min}^{-1}$ | $\text{min}^{-1}$ | $\text{min}^{-1}$ | $\text{min}^{-1}$ | – | – | – | – | – | – | $h$ |
| Fig. 5 H, I, J | 1.5e3 | 0.3 | – | – | 18 | 12 | 12<sup>a</sup> | 1 | 1.0e–8 | 1.6e–7 | 150 | 0.8 | 1.5 | 2 | 2 | – | – | – |
| Fig. 5 K, L | 1.5e3 | 0.3 | – | – | 18 | 12 | 12 | 1 | 0.25 | 1 | 150 | 0.8 | 1.5 | 2 | 1–1000 | – | – | – |
| Fig. 6 D, E<sup>b</sup> | 1.5e3 | 0.3 | 6e4 | 0.3 | 18 | 12 | 12 | 1 | 0.25 | 0 | 150 | 0.8 | 1.5 | 2 | 2 | – | – | – |
| Fig. 6 G, H<sup>c</sup> | 1.5e3 | 0.3 | 4e4 | 0.3 | 18 | 12 | 12 | 1 | 0.25 | 0 | 150 | 0.8 | 1.5 | 2 | 2 | – | – | – |
| Fig. 6 L, M<sup>d</sup> | 6e2 | 0.35 | – | – | 18 | 12 | 12 | 1 | 0.25 | 1 | 150 | 0.8 | 1.5 | 2 | 2 | 1e–4 | 6e4 | 2 |

<sup>a</sup>Experimentally determined value from (Dworkin et al., 2007)
<sup>b</sup>Overexpression of $X_R$
<sup>c</sup>Overexpression of $X_A$
<sup>d</sup>Overexpression of external repressor

Due to the fact that Antp expression takes place at the timescale of several hours to days, we can further simplify our model from (S39). In particular, we can make use of a quasi-steady state assumption (Rao and Arkin, 2003), by assuming that promoter switching due to binding and unbinding of the different Antp isoforms occurs at a much faster timescale than production and degradation of Antp. As a consequence, we can replace the stochastic gene expression rates of the two isoforms by their expected value, whereas the expectation is taken with respect to the quasi-stationary distribution of the three-state promoter model. More precisely, we have:

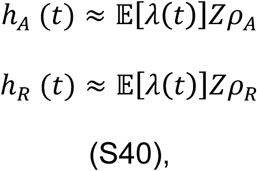

with 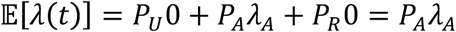 as the quasi-stationary probabilities of finding the promoter in state “U”, “A” and “R”, respectively. These probabilities can be derived from the generator matrix of the three-state promoter model, which reads:

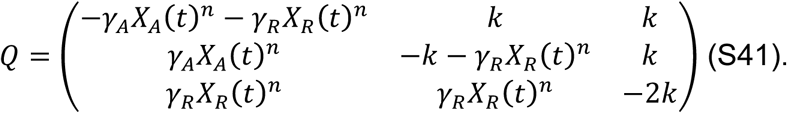

Assuming that *X_A_*(*t*) and *X_R_*(*t*) remain roughly constant on the timescale of the promoter, the quasi-stationary distribution can be determined by the null-space of Q, which is given by:

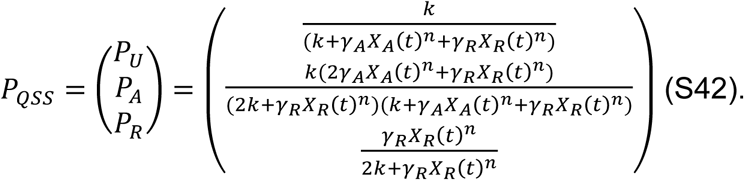

Correspondingly, the expectation of *λ*(*t*) becomes:

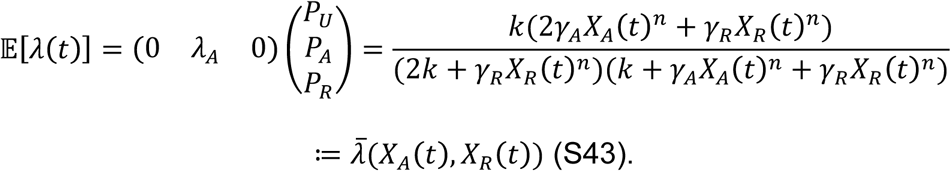

The simplified model of Antp expression can then be compactly written as two coupled birth-and-death processes:

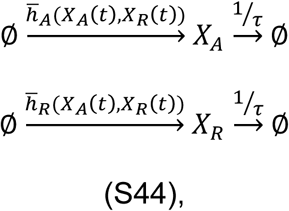

with 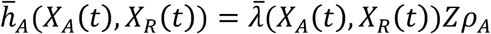 and 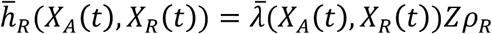.

In all our simulation studies, the circuit from (S44) was simulated using the τ-leaping algorithm (Gillespie, 2007). In case of the perturbation experiments, small modifications to the model were made. Overexpression of either of the two isoforms was reflected by changing the initial conditions of Antp. In particular, we added to the overexpressed isoform a random number of molecules drawn from a negative binomial distribution with mean *μ_0_* and squared coefficient of variation 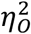 (see Table 1). To account for overexpression of an external repressor *S,* we introduced a fourth state in the promoter model, from which no expression can take place. This state is assumed to be reachable from any of the other three states at a rate 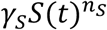 with *S*(*t*) as the concentration of the external repressor at time *t* and *n_S_* as a coefficient accounting for cooperativity in the binding of the repressor to the promoter. For simplicity, we assumed *n_S_ = n* in our case studies. To account for cell-to-cell variability in the repressor concentration, the latter was initialized randomly according to a Poisson distribution, i.e., *S*(*t*_0_)*~Pois*(*Zμ_S_*) with *μ_S_* as the average repressor abundance and *Z* as the Gamma-distributed random variable defined above. Furthermore, repressor molecules were assumed to have an average lifetime of *τ_S_*, i.e., 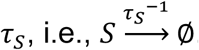.

The corresponding reaction rates of Antp expression were determined analogously to equations (S41–S43). Table 1 summarizes the parameters used for each of the simulation studies.

